# Methods for detecting PER2::LUCIFERASE bioluminescence rhythms in freely moving mice

**DOI:** 10.1101/2020.08.24.264531

**Authors:** B. Martin-Burgos, W. Wang, I. William, S. Tir, I. Mohammad, R. Javed, S. Smith, Y. Cui, C.B. Smith, V. van der Vinne, P.C. Molyneux, S.C. Miller, D. R. Weaver, T.L. Leise, M.E. Harrington

## Abstract

Circadian rhythms are driven by daily oscillations of gene expression. An important tool for studying cellular and tissue rhythms is the use of a gene reporter, such as bioluminescence from the reporter gene luciferase controlled by a rhythmically expressed gene of interest. Here we describe methods that allow measurement of bioluminescence from a freely-moving mouse housed in a standard cage. Using a LumiCycle *In Vivo* (Actimetrics), we determined conditions that allow detection of circadian rhythms of bioluminescence from the PER2 reporter, PER2::LUC, in freely behaving mice. We tested delivery of D-luciferin via a subcutaneous minipump and in the drinking water. Further, we demonstrate that a synthetic luciferase substrate, CycLuc1, can support circadian rhythms of bioluminescence, even when delivered at a lower concentration than D-luciferin. We share our analysis scripts and suggestions for further improvements in this method. This approach will be straightforward to apply to mice with tissue-specific reporters, allowing insights into responses of specific peripheral clocks to perturbations such as environmental or pharmacological manipulations.

---

Circadian rhythms arise from cycles of gene expression that occur in cells throughout the body. These cellular rhythms can be observed using a transgenic mouse with a circadian clock promoter or protein, linked to a reporter that generates signal as fluorescence or bioluminescence. The mPer2^Luc^ mouse, created in the laboratory of Joseph Takahashi (Yoo et al. 2004), has allowed many major advances in our understanding of circadian biology, largely through studies conducted using explant culture techniques. The transgene in this animal produces a fusion protein linking PERIOD2 (PER2) with firefly luciferase. Firefly luciferase, when provided with a substrate such as D-luciferin, will emit light (bioluminescence; for an overview, see (Shimomura 2012).

Bioluminescence has several advantages compared to fluorescence for long-term studies, including low phototoxicity and very low background noise (Troy et al. 2004). Studies of bioluminescence in tissue explants can reveal cellular activity, but can currently only assess isolated tissues outside their natural physiochemical environment. Additionally, in some circumstances the time of dissection for tissue preparation can reset the phase of cellular circadian clocks (Noguchi et al. 2018, Leise et al. 2018), thereby limiting the comparison of circadian phase recorded *in vitro* to the preceding phase *in vivo*. More complex interactions between tissues are best studied within a living intact animal.

To address these challenges, we and others have explored methods for recording bioluminescence *in vivo*. One approach is to use anesthetized animals, detecting bioluminescence using a camera-based system. This approach has been used to assess the phase of peripheral tissue rhythms (Tahara et al. 2012) but has the unavoidable disadvantage of repeated general anesthesia. Anesthesia has been shown to disrupt circadian rhythms in body temperature and in rest/activity cycles, and may inhibit photic entrainment of the circadian clock (Poulsen et al. 2018). The method also requires bioluminescence assessments at six to nine time points, each requiring experimenter presence, over 24 to 36 hours for a single circadian phase assessment. A second approach is to record bioluminescence from freely behaving mice, detecting light emission using sensitive photomultiplier tubes (PMTs). Here we build on the pioneering work from the Schibler laboratory (Saini et al. 2013) that demonstrated the feasibility of recording light emitted from a mouse in a specialized housing chamber, the “RT Biolumicorder” (LESA Technology). In their work, rhythmic signal was detected from bioluminescent reporters directed to the liver using viral transduction in hairless mice, as well as in m*Per2^Luc^* C57BL/6 mice with fur removed from the dorsal region. In addition to establishing feasibility of the method, the system was used to control food availability and provide lighting regimens for studies of entrainment. Some areas of further research are to develop a system that allows housing in standard cages, to test nonsurgical methods for delivery of D-luciferin, and to test synthetic luciferins that may offer increased sensitivity (Evans et al. 2014).

We present here results from studies of PER2::LUC bioluminescence in C57BL/6 mice, recorded in standard housing cages within the “LumiCycle *In Vivo*” system (designed for these experiments by Actimetrics Inc.). We compare two methods of delivery for the substrate D-luciferin: delivery through drinking water, and using subcutaneous osmotic mini-pumps. Additionally, we assessed whether a synthetic luciferin, CycLuc1 (Evans et al. 2014), can offer advantages as a substrate in these experiments. In the course of this work we encountered several methodological challenges and addressed how they may be overcome. We believe that these methods could be widely adopted and could also be improved through further innovation. Ultimately, we predict the field of circadian biology will benefit from expanded use of unrestrained, *in vivo*, bioluminescence recordings.

## Methods

### The LumiCycle *In Vivo*

Bioluminescence was measured using the “LumiCycle *In Vivo*” system, designed by David Ferster (Actimetrics). The system can record luminescent emissions from freely behaving animals housed in a standard mouse cage placed within a light-tight chamber (Supplementary Figure S1).

Each unit contained two PMTs (Hamamatsu H8259-01, selected for low dark counts and positioned for sensitivity across the cage area), and was equipped with programmable LED lights, and a high-capacity internal fan that provided 15 air changes per hour. A programmable shutter periodically blocked the PMTs and allowed the collection of dark counts. In the studies reported here, dark counts were measured every 15 minutes by closing the shutter for 1 minute. Each dark-count value was subtracted from the counts recorded during the subsequent 14 minutes to obtain the background-corrected count values. This was necessary to remove the effect of external temperature fluctuations on PMT noise. Prior to this correction we observed noise highly correlated to temperature (R^2^= 0.96) and less so with humidity (R^2^=0.30). One degree Celsius of chamber ambient temperature increase was associated with an increase of circa 35 counts/s. A chamber history for all recording units was collected, to monitor temperature, relative humidity, and light measures within each unit. Animal welfare was checked once a day at varied times using an infrared viewer (Carson OPMOD DNV 1.0), or goggles (Pulsar Edge Night Vision Goggles PL75095).

### Animals

For studies conducted at Smith College, animals were bred in-house from founders obtained from Jackson Labs (JAX #006852). Homozygous mPer2^Luc/Luc^ male and female mice were included in this study (Yoo et al. 2004), ranging in age from 4.5 to 8 months. We chose homozygous mice in order to maximize luciferase expression.

IVIS imaging studies were conducted at the University of Massachusetts Medical School, and employed either FVB/NJ mice purchased from Jackson Laboratories and given intravenous injection of AAV9-CMV-WTluc2 as previously described (Mofford et al. 2015) or, in separate studies, mice with a single copy of the Per2::LucSV allele (Welsh et al. 2004, Yoo et al. 2017). The latter mice are referred to as mPer2^LucSV/+^ to distinguish them from the homozygous mice used elsewhere. All animal studies were approved by the Institutional Animal Care and Use Committees of the appropriate institution.

### Delivery of substrates by subcutaneous osmotic mini-pump

A delivery method that can provide continuous dosing is to use a subcutaneous osmotic mini-pump. Prior studies have reported mixed results with similar methods (Gross et al. 2007, Saini et al. 2013, Hamada et al. 2016). We determined that 100 mM was close to the limit of solubility for D-luciferin in our conditions and therefore tested this concentration in both 7-day and 14-day Alzet osmotic minipumps (models 1007D and 1002, respectively) to vary the rate of delivery (0.5 and 0.25 ul/h). 100 mM D-luciferin was prepared in sterile 0.1 M sodium phosphate buffer (PB) and mixed on an Eppendorf ThermoMixer (Thermo Fisher Scientific, Waltham, MA) at 37° C for two minutes @ 1500 RPM. Following surgery to implant an osmotic mini-pump subcutaneously we allowed 3 days recovery before initiating bioluminescence recording (see Supplementary Methods for details).

A synthetic luciferin, CycLuc1, has been shown in prior studies to offer better distribution throughout the body and greater light emissions at lower concentrations (Evans et al. 2014). We tested whether this synthetic luciferin could provide comparable or superior results to D-luciferin when delivered via an osmotic pump. CycLuc1 (produced in the lab of Dr. Stephen Miller, University of Massachusetts Medical School, Worcester, Massachusetts) was prepared in a 5 mM solution in sterile 0.1 M phosphate-buffered saline (PBS) and was mixed as for D-luciferin. Both 7-day and 14-day pumps were tested.

The bioluminescence data was collected until day 9 for 7-day pumps and until day 11 or 12 for 14-day pumps. After removal, any residual volume remaining in pumps was measured to check against the expected rate of delivery, and any changes in the appearance of the solution were noted. Data files were truncated if necessary to remove measures taken after calculated pump depletion.

Each animal underwent two minipump trials using the procedures described above. The two pump implantation surgeries were separated by 28 days and within each group of surgeries pump volumes were selected in a counterbalanced design. Each animal received the same substrate on both occasions, with only the pump type (7-day or 14-day) differing.

### Delivery of D-luciferin by drinking water

A technically simple manner of substrate delivery is to provide the compound in the drinking water. A prior study suggested this approach of substrate delivery would be feasible (Hiler et al. 2006). This approach depends on the availability of a reasonably inexpensive substrate in appropriate amounts, solubility, acceptability by the mouse, and low toxicity. For our initial experiments, D-luciferin, potassium salt (LUCK-1G, GoldBio, St Louis, MO) was selected. Major advantages of this approach are the potential to conduct long-term studies with fresh solutions provided at regular intervals, and the avoidance of surgical procedures. A concern with administration of substrate in the drinking water is the possibility that the rhythmic behavior of drinking would lead to rhythmicity in substrate ingestion and availability, thus producing bioluminescence rhythms that reflect drinking behavior rather than reporter gene expression. This, however, is best addressed using mice with peak reporter expression at a different phase from peak drinking behavior, and is not addressed in the experiments reported here.

To study dose-response to D-luciferin through drinking and reversibility, each mouse was tested for approximately 3 days at each concentration. We tested 10 mM, 0 mM, 1 mM, and 5 mM of D-luciferin, in that order. Solutions were made by dissolving powdered D-luciferin in water in 50 mL conical tubes, vortexing 1-2 minutes to dissolve particulate and then protected from light. Experiments were thereafter conducted in constant darkness. Drinking was monitored by measuring residual volume and all doses were equally tolerated by the mice tested. An additional 3 day experiment with 0 mM D-luciferin was conducted 5 weeks later, to test baseline levels when it was certain that no substrate remained within the animal after prior exposure. On the first day of this experiment as well as immediately prior to the recording conducted 5 weeks later, animals were anesthetized briefly (circa 5 min) in an induction chamber at 3% isoflurane followed by placement in a nose cone to allow shaving of back and stomach (see Supplementary Figure S3).

Animals were put into the LumiCycle *In Vivo* unit at the end of the light portion of the 12:12 LD cycle and were thereafter recorded in constant darkness. The bottle of D-luciferin was changed to the new dose every 3 days between CT 6 and CT 11. Mice drank on average 3.9 mL/ day.

To compare D-luciferin and CycLuc1 delivered in drinking water, four mice were tested using an ABA design with 0.1 mM D-luciferin, 0.1 mM CycLuc1, and a second treatment of 0.1 mM D-luciferin. Treatments were given for approximately 3 days each, with approximately 1 day with plain water available in between each treatment. Mice were given Cycluc1 for 4 days, due to a circa-1 day equipment failure during this experimental run. Bioluminescence was measured in DD using the LumiCycle *In Vivo* unit, and volume consumed was monitored by noting initial and final water volumes.

### Assessment of the source of bioluminescence by IVIS imaging

mPer2^LucSV/+^ mice were allowed *ad libitum* access to drinking water containing D-luciferin (2 mM, n=10 or 0.1 mM, n=6) or CycLuc1 (0.1 mM, n=10) for at least 12 hours before study. Another group of animals (n=7) had water available and received D-luciferin by injection (0.1 mL of 10 mM, i.p.). Animals were studied at clock times at which elevated bioluminescence was expected based on our previous studies (ZT15-21, van der Vinne et al., 2018). Mice were anesthetized with isoflurane, removed briefly to be shaved, and images were captured of ventral and dorsal views using Living Image software on an In Vivo Imaging System (IVIS-100, Caliper, now Perkin Elmer) in the UMass Medical School Small Animal Imaging Core Facility. After external imaging the animals were euthanized by decapitation while under anesthesia, and organs were dissected. Images of the dissected tissues were captured to assess the distribution of light coming from specific anatomical areas. Quantitative analysis of bioluminescence was performed using Living Image 4.4 software.

To assess the distribution of bioluminescence among internal organs, values for each tissue were expressed as a percentage of the total signal from all identified anatomical areas. To accommodate inconsistencies in dissections, the signal from uterus, perigonadal fat and any dissected intra-abdominal adipose tissue were summed in females to a single value described called “fat”. In males, epididymal adipose tissue and any dissected intra-abdominal adipose tissue were summed to give a single value “fat”. Several tissues did not contribute substantially (<4%) to total signal and are lumped together as “other”. These tissues (and the number of observations leading to their assignment to this group) are: esophagus (16), stomach (44) lung (39), heart (40), thymus (32), spleen (41), testes (19), seminal vesicles (16), urinary bladder (24) brain (9), and head (40). Values from the remaining carcass post-dissection were labeled “body”.

### Data analysis

All data were analyzed using RStudio, with the exception of the Bayesian MCMC analysis, which was run in MATLAB. Data files and R code are posted on Open Science Framework (Link for peer review purposes: https://osf.io/p3qke/?view_only=41db92d90ade49ea8c2db2537532ed11). A discrete wavelet transform (DWT) was applied to each time series to detrend and to calculate the rhythm phase using the wmtsa R package (https://cran.r-project.org/web/packages/wmtsa/index.html), as described in (Leise and Harrington 2011, Leise 2017). We applied the S12 filter on 15-min median binned data; medians were used to reduce the effect of large outliers. Data before the first trough and after the last trough were discarded to avoid edge effects.

The signal-to-noise ratio (SNR) in decibels was calculated for 2-day windows, starting at the first trough (dropping most of the first day of recording), as 10 times the log10 of the ratio of the sum of squares of DWT circadian component D6 to the sum of squares of all DWT components at least 2 levels below the circadian scale (D1-D4). The level immediately below circadian (D5) contains an inseparable mix of circadian and noise energy, so was discarded. An SNR value above 0 indicates the signal is stronger than the noise (ratio was greater than 1). The SNR does not statistically test whether there is a significant circadian rhythm present, although a high SNR value strongly suggests it. A negative SNR value can mean either a rhythm swamped by noise or no rhythm (ratio of signal to noise energy was less than 1).

The circadian energy is the sum of squares of the DWT circadian component D6 divided by the sum of squares of all DWT components D1-D6, yielding the proportion of total energy that is at the circadian scale, for the same window as the SNR. This proportion does not indicate whether there is a significant circadian rhythm present, although a value above 0.60 strongly suggests it.

As a measure of uncertainty in whether the period can be determined from the data, we ran a Bayesian statistical analysis via MCMC sinusoid fitting, as described in (Cohen et al. 2012), for 4-day windows, dropping the first 24h of recording. The procedure generates a distribution of period estimates, where the 95% credible interval (CI) is the interval of most likely period values. A mean period estimate can also be obtained from this distribution. The CI provides evidence for whether the period of a time series can be reliably measured. The wider the CI, the less credible it is that the time series has a measurable circadian period. Only 14-day pump records were used for this computation, to have sufficient length.

To assess the statistical significance of circadian rhythms in the data, JTK and Lomb-Scargle q-values (false-discovery-rate adjusted p-values) were calculated using the MetaCycle R package (Wu et al. 2016), applied to 3-day windows, dropping the first 24h of the recording and using 90-min median binning to reduce correlated high-frequency noise and outliers.

DWT peak times are the peaks in the circadian signal component of the bioluminescence obtained using the discrete wavelet transform. Time relative to projected lights indicate the DWT peak times in relation to the time of lights on (6 am) in the colony room where animals were housed prior to the *in vivo* bioluminescence experiments.

Further experimental details are provided in the supplementary methods.

## Results

As in many previous reports (Saini et al. 2013, van der Vinne et al. 2018, Crosby et al. 2019), we were able to detect circadian rhythms of bioluminescence *in vivo* from mice carrying the mPer2^Luc^ transgene (see Figure 1). Given that the reporter signal arises from multiple tissues (see below), this suggests that the tissues we are detecting express rhythms with some degree of synchrony even during housing in constant darkness.

**Figure 1.**
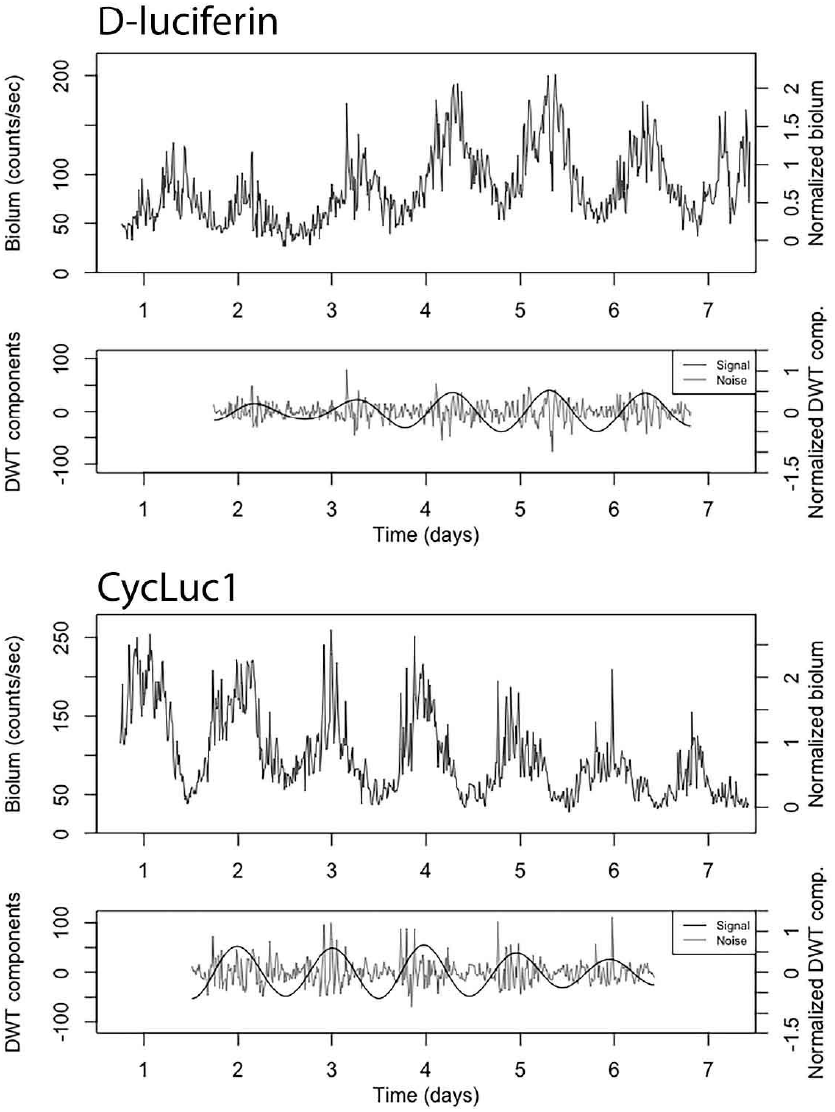
Example of bioluminescence data with 14-day D-luciferin pump (0.25 uL/h of 100 mM D-Luciferin), or 14-day CycLuc1 pump (0.25 uL/h of 5 mM CycLuc1). Top graph for each example: 15-minute median binned trace with counts/sec scale on the left and normalized scale on the right (subtract first percentile then divide by median, so min is mapped to zero and median mapped to 1; using first percentile reduces the effect of outliers). Bottom graph for each example: For the signal-to-noise ratio analysis, the DWT-calculated circadian component D6 is treated as the signal and the summed components D1-D4 are treated as the noise. The data before the first trough and after the last trough are discarded to avoid edge effects.

We first investigated bioluminescence rhythms in mice receiving D-luciferin continuously using an osmotic minipump (Figure 1). A 14-day pump delivers a smaller volume per h than a 7-day pump (0.25 uL/h vs 0.5 ul/h, respectively) yet both pumps are limited to a 100ul reservoir. We asked if this reservoir, with 100 mM D-luciferin, was able to lead to a detectable rhythm in bioluminescence, in a small sample of 4 mice. This experiment also tested the impact of holding D-luciferin solution at body temperature for up to 11-12 days. Table 1 shows the results. One animal was noted to have unusually rapid fur regrowth and showed progressive signal loss (see Supplementary Figure S4C). Both rates of D-luciferin delivery led to detectable circadian rhythms in bioluminescence.

**Table 1.** Summary of key measures of rhythms recorded from mice with osmotic pumps delivering substrate. See Methods for details. Peak time is given in hours with 0=midnight, using the peak nearest the day 2 mark. Prior housing room LD cycle had lights on 0600-1800 h. Median bioluminescence is for the first 3 days of recording.

| ID | Sex | Substrate | Pump | Median Biolum | Peak time (h) | Time relative to projected lights on (h) | SNR | Circ. energy | CI width | Mean period | JTK q-val | LS q-val |
| --- | --- | --- | --- | --- | --- | --- | --- | --- | --- | --- | --- | --- |
| 82 | M | D-luciferin | 14-d | 62 | 6.5 | 24.5 | -1.0 | 0.50 | 2.3 h | 23.2 h | 0.01 | 0.062 |
|  |  |  | 7-d | 157 | 6.8 | 24.8 | -3.2 | 0.36 | -- | -- | <0.001 | <0.001 |
| 83 | M | D-luciferin | 14-d | 20 | 4.0 | 22.0 | -9.9 | 0.12 | NA | NA | 0.1 | 0.262 |
|  |  |  | 7-d | 72 | 3.5 | 21.5 | 1.0 | 0.49 | -- | -- | <0.001 | <0.001 |
| 84 | F | D-luciferin | 14-d | 42 | 0.3 | 18.3 | 4.8 | 0.76 | 0.9 h | 24.5 h | <0.001 | <0.001 |
|  |  |  | 7-d | 46 | 1.3 | 19.3 | 1.8 | 0.62 | -- | -- | <0.001 | <0.001 |
| 85 | F | D-luciferin | 14-d | 22 | 1.8 | 19.8 | -0.9 | 0.40 | 1.4 h | 23.9 h | 0.002 | 0.009 |
|  |  |  | 7-d | 30 | 1.5 | 19.5 | 4.4 | 0.63 | -- | -- | <0.001 | <0.001 |
| 90 | M | CycLuc1 | 14-d | 76 | 6.8 | 24.8 | -1.2 | 0.44 | 1.5 h | 24.0 h | 0.002 | 0.009 |
|  |  |  | 7-d | 140 | 0.8 | 18.8 | 1.7 | 0.61 | -- | -- | <0.001 | <0.001 |
| 91 | M | CycLuc1 | 14-d | 237 | 5.8 | 23.8 | -3.0 | 0.38 | 1.5 h | 24.9 h | <0.001 | <0.001 |
|  |  |  | 7-d | 467 | 6.3 | 24.3 | -5.0 | 0.28 | -- | -- | <0.001 | <0.001 |
| 92 | F | CycLuc1 | 14-d | 92 | 1.3 | 19.3 | 2.6 | 0.67 | 1.2 h | 24.5 h | <0.001 | <0.001 |
|  |  |  | 7-d | 173 | 1.0 | 19.0 | 5.3 | 0.80 | -- | -- | <0.001 | <0.001 |
| 93 | F | CycLuc1 | 14-d | 108 | 0.0 | 18.0 | 3.1 | 0.69 | 0.9 h | 23.6 h | <0.001 | <0.001 |
|  |  |  | 7-d | 106 | 0.3 | 18.3 | 1.4 | 0.62 | -- | -- | <0.001 | <0.001 |

In addition, we asked if the synthetic luciferase substrate, CycLuc1, can be effective at concentrations lower than that of D-luciferin, as previously seen in studies with acute injection of the substrates (Evans et al. 2014). We tested 5 mM CycLuc1 in 7-day and 14-day osmotic minipumps, and again we detected circadian rhythms in bioluminescence (Figure 1). Traces for all 16 recordings (7- and 14-day pumps, both substrates) are given in Supplementary Figure S4.

Table 1 provides a summary of the analysis of the osmotic minipump data. All but one animal had bioluminescence profiles that were significantly rhythmic according to both JTK and Lomb-Scargle tests. The exception is the one mouse with a 14-day D-luciferin recording mentioned above, for which hair regrowth was an issue. The Bayesian statistical analysis supports this assessment, yielding mean periods near 24 h and reasonable CI widths, given the shortness of the time series, again with the exception of the single problematic 14-day D-luciferin recording. The signal-to-noise ratio (SNR) varied, reflecting the high-amplitude noise present in many of these recordings; noise amplitude tended to increase greatly near signal peaks, as shown in Figures 1 and 2. While SNR values above 2 tend to allow greater precision in period and phase estimates, the analysis here suggests that even when noise levels are relatively high, the recordings are statistically rhythmic with measurable period and phase (though keeping in mind the uncertainty indicated by the Bayesian analysis). The circadian energy provides an alternative measure to the SNR for assessing signal strength. This measure largely agrees with the SNR results -- strong circadian signals tend to have SNR above 1.5 and circadian energy above 0.6.

**Figure 2.**
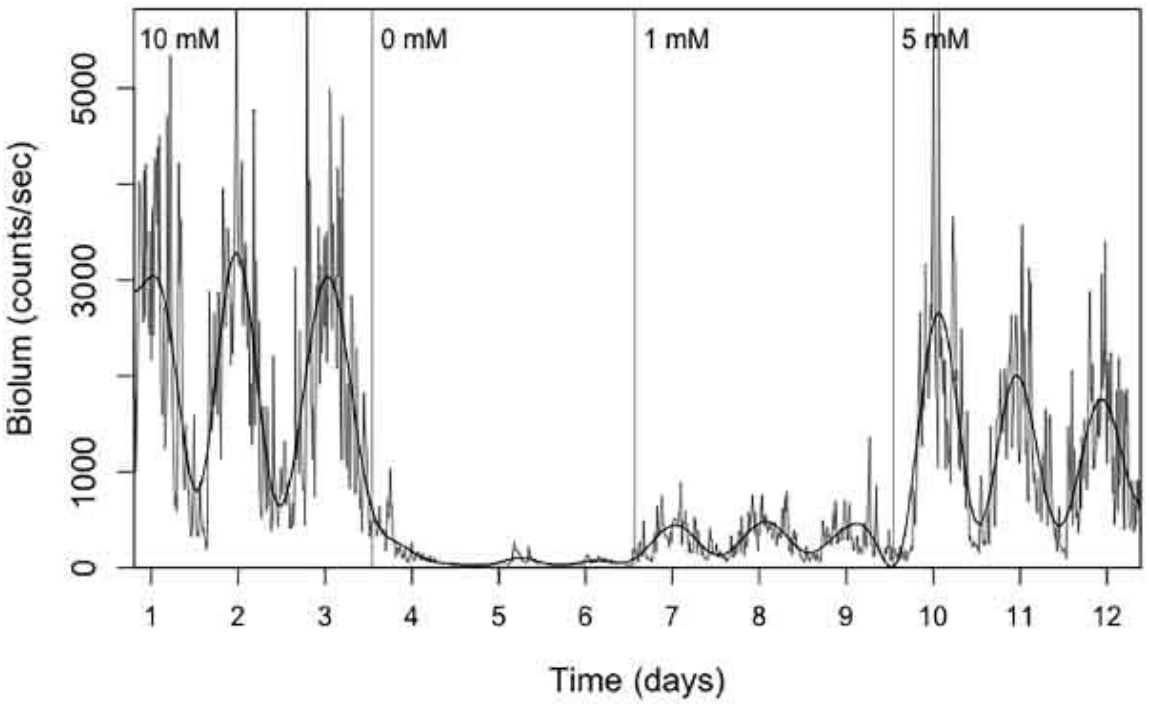
Example of bioluminescence data with D-luciferin delivered via drinking water, using a sequence of doses as indicated. Vertical lines indicate when substrate solutions were changed.

Next, we asked if we could measure rhythmic bioluminescence using delivery of substrate in the drinking water, and if the signal varied with the concentration of D-luciferin. As shown in Figure 2, we saw an increased median level of bioluminescence with increasing concentration of D-luciferin in the drinking water across the 4 mice tested, but the SNR was not reliably altered with dose. The 0 mM condition immediately following the 10 mM concentration allowed us to observe the duration of wash-out (loss of signal following substrate removal). Also shown in Figure 2, residual signal was observed for variable amounts of time after D-luciferin (10 mM) withdrawal. Notably an additional 0 mM condition that was not immediately preceded by D-luciferin administration yielded a true baseline, with low levels of bioluminescence (with negative SNR and non-significant JTK and Lomb-Scargle q-values). All drinking water bioluminescence recordings are shown in Supplemental Figure S5.

We then asked if delivery of the synthetic substrate CycLuc1 in the drinking water would produce robust bioluminescence signal at lower concentrations of substrate, as previously reported (Evans et al. 2014). As shown in Figure 3, CycLuc1 treatment (0.1 mM) produced a stronger bioluminescence signal than the same concentration of D-luciferin in each of 4 animals. We found no consistent difference in the amount mice drank across conditions (Supplemental Figure S6).

**Figure 3.**
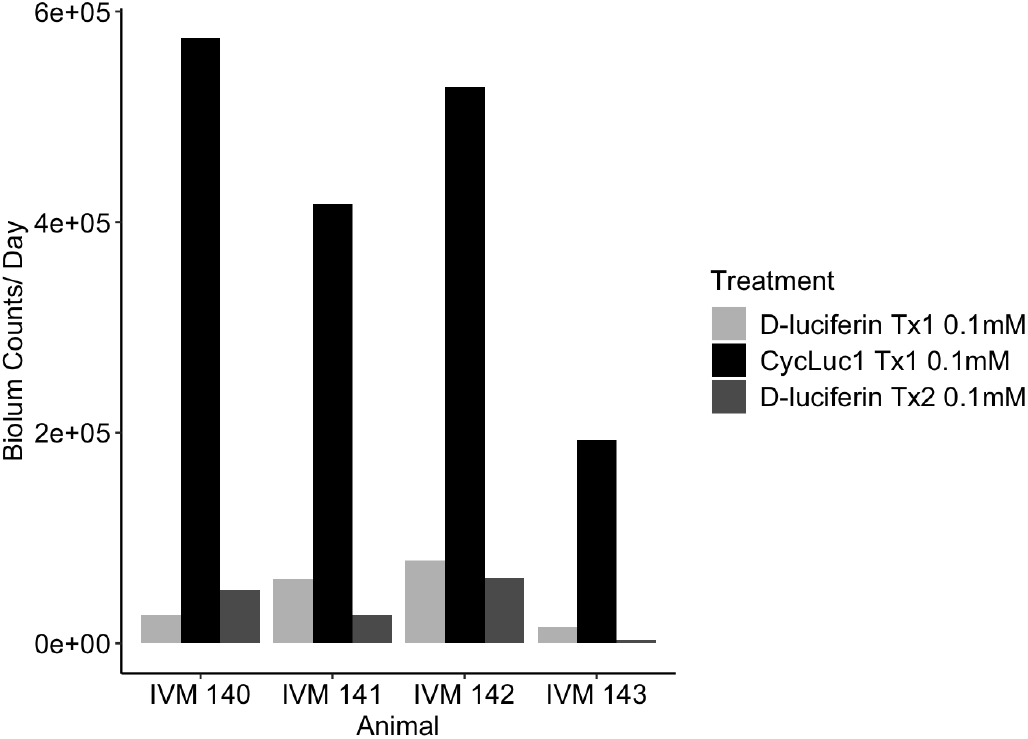
Bioluminescence counts per day were calculated for each treatment by dividing the total sum of counts by the number of days. Treatments were 0.1mM D-Luciferin, 0.1mM CycLuc1, and 0.1mM D-Luciferin, and were 3 days in length with a day of plain water administered between treatments.

Bioluminescent rhythm damping (as seen e.g. in Figure S4G) might be attributable to fur regrowth in these black mice, and/or substrate degradation over time. We incubated aqueous solutions of D-luciferin and CycLuc1 for stability for multiple days at body temperature to determine their relative stability. Prolonged incubation of D-luciferin at 37 °C resulted in substantial oxidation of D-luciferin to dehydroluciferin as judged by LC-MS (see Supplementary Methods). After 14 days, ~50% of the D-luciferin had oxidized (Figure 4). Some loss of D-luciferin by racemization to L-luciferin is also likely (Nakajima et al. 2020). Dehydroluciferin is not a luminogenic substrate for firefly luciferase, and is a much more potent inhibitor than L-luciferin (da Silva and da Silva 2011). The ~6-fold decrease in bioluminescent signal observed when using this solution for in vivo imaging may thus reflect luciferase inhibition by dehydroluciferin as well as a reduction in the D-luciferin dose. By contrast, no formation of a dehydroluciferin product was noted for CycLuc1 at 37 °C or for either substrate incubated at 4 °C, and the *in vivo* imaging results were comparable to a freshly-prepared solution (Supplemental Figure S7).

**Figure 4.**
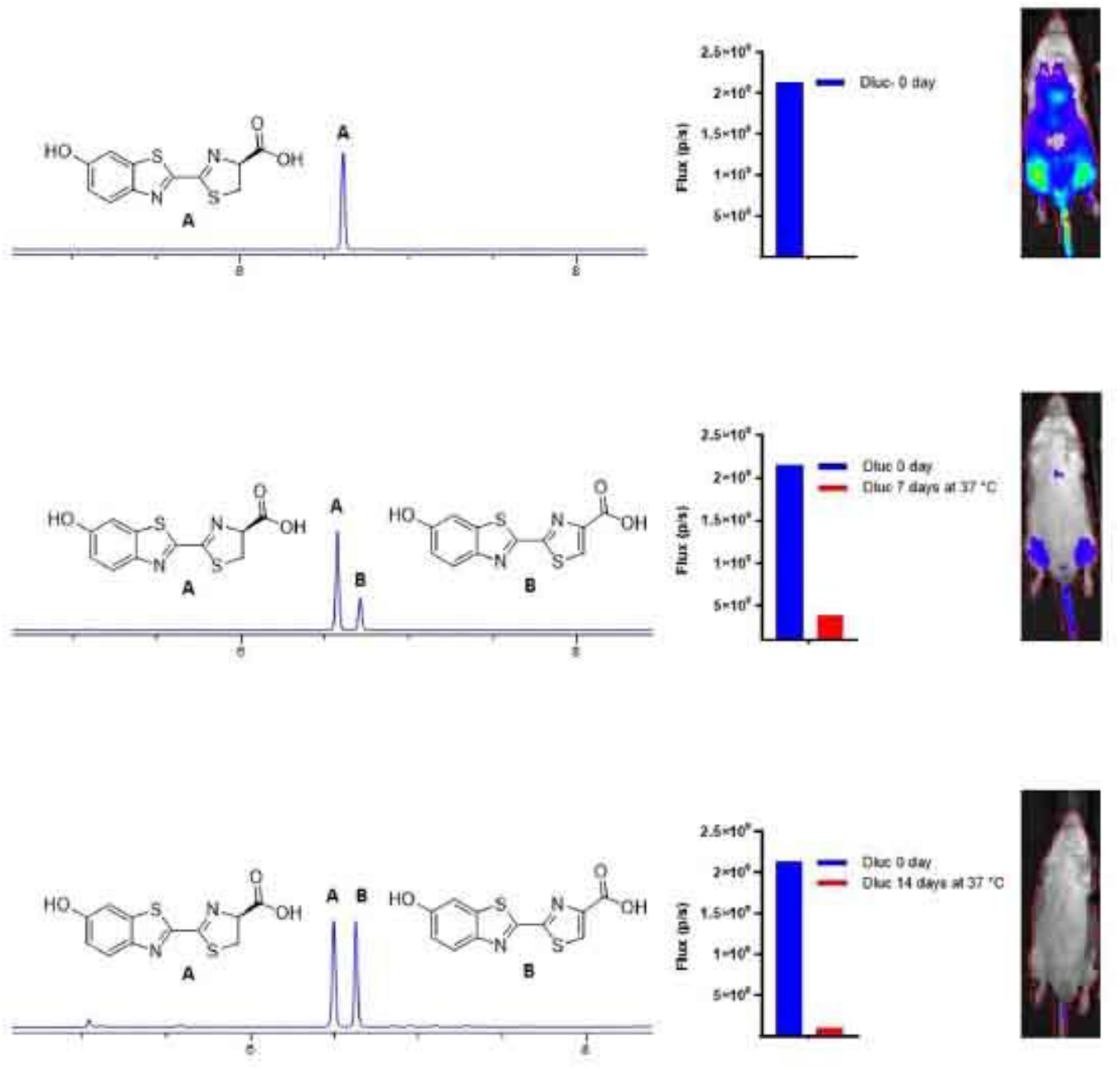
Evaluation of D-luciferin after incubation at body temperature. LC/MS and *in vivo* bioluminescence imaging with D-luciferin incubated in aqueous buffer at 37 °C for 0, 7, or 14 days.

One shortcoming of assessing bioluminescence rhythms using the LumiCycle *In Vivo* is that this method does not provide information on the anatomical source of bioluminescence. Previous studies of Per2::Luc in mice conducted with bioluminescence imaging have identified the liver, kidneys, and submandibular gland as the primary sources of ‘signal’ from anesthetized mice (Tahara et al. 2012). Notably, in each of these studies, the substrate was provided by intraperitoneal injection. Thus, we performed IVIS imaging studies to assess the sites of origin of bioluminescence in mPer2^LucSV/+^ mice receiving substrate in the drinking water.

Mice drinking D-luciferin (2 mM or 0.1 mM) or CycLuc1 (0.1 mM) had readily detectable bioluminescence signal, located predominantly in the upper abdomen (ventral view) or from the area on the dorsal surface corresponding to the abdomen (dorsal view) (see Figure 5). These regions contributed over 75% of photons within the region of interest containing the entire animal. The anatomical pattern of labeling did not appear related to the treatment group. Bioluminescence (at much lower levels) was observed in a variable pattern in extra-abdominal sites, including hands, feet, mouth, throat (presumably submandibular gland) and where urine leaves the body (vulva in females, penis in males), and on the dorsal side, upper back and rump. Mice receiving D-luciferin by injection had a similar distribution, including the preponderance of labeling coming from the abdomen in both ventral and dorsal views. We compared the log of counts from either ventral or dorsal views across conditions and sex using ANOVA. We found a significant effect of treatment in each view (Ventral: F(3,25)=107.21, p<.001; Dorsal: F(3,20)=152.2, p<.001). Tukey contrasts showed bioluminescence was greatest in mice receiving D-luciferin by injection (0.1 mL of 10 mM), followed by the two treatments CycLuc1 (0.1mM) and D-luciferin (2 mM) (no significant statistical difference between these) and finally by D-luciferin (0.1 mM). When considering the ventral view, we saw no significant effect of sex (F(1,25)=0.002, p=.96) but for the dorsal views we found a weak sex effect (F(1,20)=4.59, p=.0446; signal trended higher in males).

**Figure 5.**
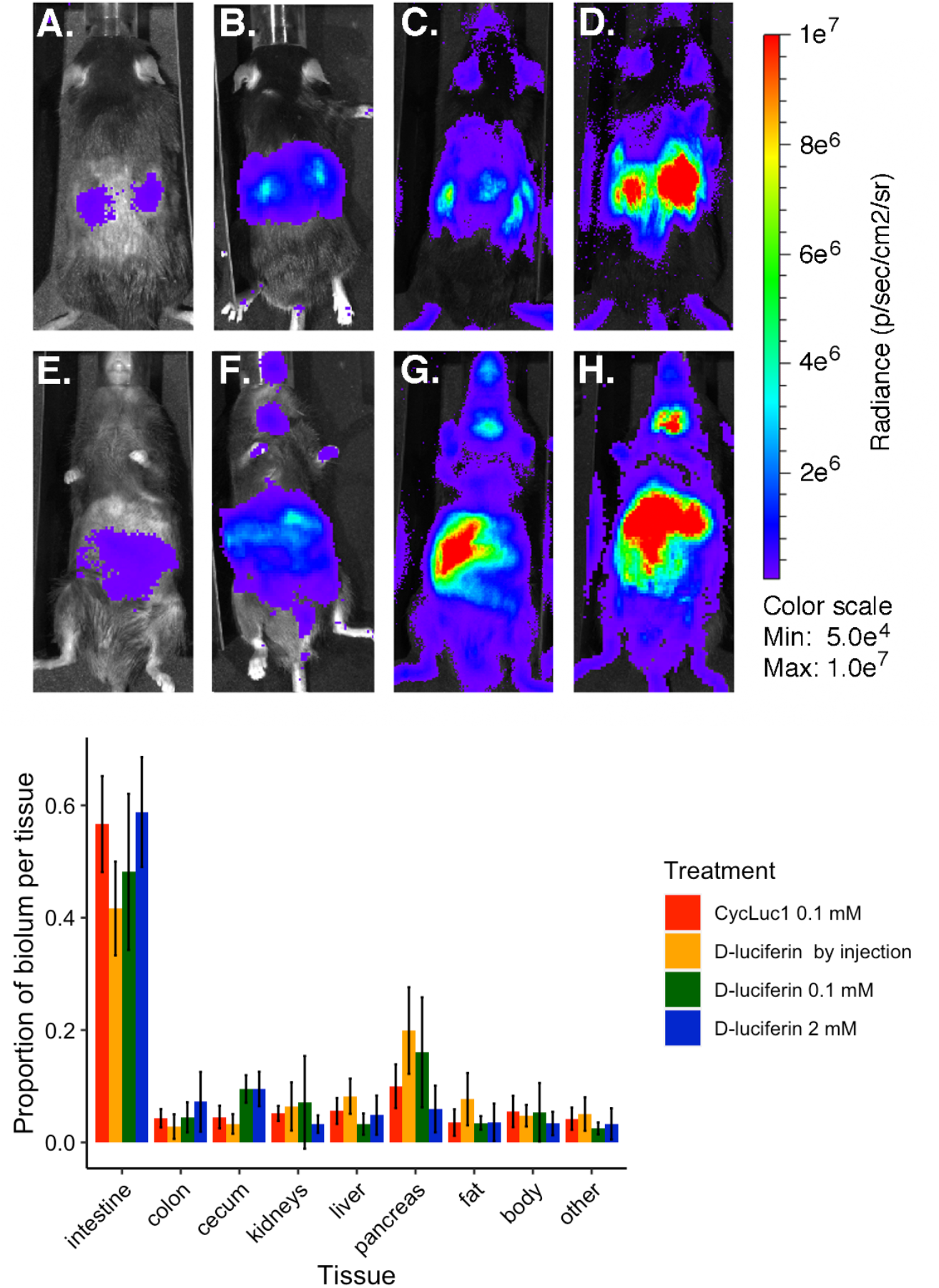
IVIS imaging to assess the source of bioluminescence in Per2^LucSV/+^ mice. <u>Upper panel:</u> Images show paired dorsal (A-D) and ventral (E-H) images from a representative male mouse from each treatment group. Mice were imaged after drinking 0.1 mM D-luciferin (A: dorsal view, E: ventral view), 2 mM D-luciferin (B: dorsal view, F: ventral view) or CycLuc1 (0.1 mM; C: dorsal view, G: ventral view), or after injection of 0.1mL D-luciferin (10mM; D: dorsal view, H: ventral view). All images are adjusted to the bioluminescence scale bar shown at right. <u>Lower panel:</u> Proportion of total bioluminescence in each tissue assessed by IVIS imaging after dissection is summarized by mean +/− SD.

To further localize the source of bioluminescence signal, mice were euthanized by cervical dislocation while anesthetized with isoflurane, and tissues were rapidly dissected, spread out on paper, and images were captured. Luciferase activity is dependent upon temperature, ATP and oxygen, all of which would plummet rapidly after euthanasia. Assuming that the decline in these constituents occurs at an equivalent rate in different tissues, we can infer the relative strength of signals during *in vivo* imaging, based on the pattern seen in dissected tissues. Bioluminescence signal was strongest from tissues of the gastrointestinal tract in all animals. The intestine, cecum and colon were highly but heterogeneously labeled, and together accounted for ~65% of counts. The pancreas and mesenteric tissue were not readily separated and had similar signal intensity, so this was imaged as a single ROI (labeled “pancreas” in lower panel, Figure 5); this tissue contributed ~ 10% of signal. Liver and kidney each contributed ~ 6%. The only other regions that consistently had detectable signal were the remainder of the body (from which tissues had been removed), and adipose tissue (especially perigonadal fat), which also averaged ~ 4-6% of light output. Given these findings, we suspect that the presence of higher bioluminescence in the kidney region on the dorsal view of intact mice might be due, in large part, to the spinal column blocking bioluminescence arising from the abdomen.

Considering the total light output from these tissues, combined with their anatomical location (and likely physical obstacles to bioluminescence detection *in vivo*), it appears that the major contributors to bioluminescence signal detected from ambulatory mice are gastrointestinal tissues (intestine, cecum and colon, and pancreas), and kidney (by virtue of its dorsal location). Unexpectedly, liver was not a major contributor to the bioluminescence signal. We did not observe an obvious difference in source of bioluminescence depending on the substrate or route of administration (see lower panel, Figure 5).

## Discussion

These studies extend prior work that established the feasibility of recording gene expression rhythms *in vivo* with bioluminescence. We have chosen to conduct initial studies with the most widely used animal model for circadian bioluminescence currently available, the m*Per2^Luc/Luc^* mouse with the reporter PER2::LUC. We used mice with two copies of the reporter allele to enhance signal output. Thus, success with this animal model likely sets the mark for the most we can achieve, with less light expected from animals with bioluminescence directed to specific organs. On the other hand, PER2::LUC may offer a noisier rhythmic signal than organ-specific reporters, because it is reporting from diverse sources of signal. These diverse sources must be somewhat coordinated, to detect circadian rhythms at the whole-animal level as reported here. This is consistent with data from reports assessing bioluminescence rhythms in anesthetized mice by IVIS imaging, in which rhythms of PER2::LUC bioluminescence were relatively synchronous across the tissues assessed (Tahara et al. 2012, van der Vinne et al. 2018).

Our results indicate that luciferin substrates are effective for producing detectable bioluminescence rhythms when delivered either by drinking water or by osmotic minipump. The possibility that the daily drinking rhythm altered the phase and/or amplitude of the bioluminescence rhythm could not be excluded in the current study. Future studies should be conducted using mice with a circadian reporter with peak expression at a phase not coincident with the major time of voluntary water intake. In the studies here, phase was rather variable, perhaps not unexpected when using mice with a reporter expressed throughout the body.

The synthetic luciferin, CycLuc1, offers advantages in terms of signal detection and stability. CycLuc1 has a lower Km for firefly luciferase than D-luciferin, and can be used effectively at lower doses (Evans et al. 2014). The ability of this substrate to be used at lower concentrations and/or delivered at slower rates, could potentially overcome some of the limitations imposed by the small volume of minipump reservoirs. Use of CycLuc1 might offer the possibility for longer-term experiments, with multiple days of recording providing estimates of circadian period with increased precision (Cohen et al. 2012).

Another consideration for long-term studies using implanted osmotic pumps is the aqueous stability of the luciferin substrate at body temperature. Oxidation of a luciferin to its respective dehydroluciferin reduces the luciferin concentration and also introduces a potent luciferase inhibitor.

We explored several different analysis methods for assessing the strength of the rhythmic signal in the *in vivo* bioluminescence data. The signal-to-noise ratio can be useful for measuring improvements in reducing the high frequency noise to create a clearer circadian signal, as can the circadian energy measure. These two measures provide evidence toward whether the period and phase can be reliably estimated, but are not direct tests of rhythmicity. Similarly, the Bayesian statistical analysis provides a measure of how reliable the period estimate is, but is not a rhythmicity test. Finding an effective rhythmicity test proved to be challenging, due to the correlated nature of the high frequency noise which caused some tests to give spurious results. The two statistical tests that appeared to work most reliably were JTK_CYCLE and Lomb-Scargle, as implemented in MetaCycle (Wu et al. 2016), using 90-min bins to avoid the problematic effects of the high frequency noise. The general conclusion from the various methods is that the *in vivo* methods are generating data with clear circadian rhythms for which period and phase can be estimated with reasonable precision, analogous to that for other data with a similar number of cycles.

Our DWT peak pump results (Table 1) suggested a possible sex difference in the timing of the measured rhythms, yet our sample size did not allow firm conclusions. To explore potential DWT bioluminescence peak differences between whole body male and female mice expressing PER2::LUC, we applied our DWT analysis on a combined dataset including all osmotic minipump treatments presented in this study, and a group of controls from another recently published paper (van der Vinne et al. 2020) that studied metabolic consequences and PER2::LUC bioluminescence in mice lacking a functional central clock (*Vgat-Cre^+^ Bmal1^fl/f^,* experimental group). The control animals from this study (No Cre *Bmal1^fl/f^ Per2^LucSV/+^*) are another translational reporter from the *Per2* locus, similar to the m*Per2^Luc/Luc^* animals used for ambulatory recordings in the current study. Bioluminescence data were obtained in our lab using the same equipment and methods presented in this paper, with CycLuc1 substrate delivered at a 0.25 uL/h rate in 14-day pumps. The DWT peak analysis (Figure 6) showed more variation in DWT peak times in males than females. We did not apply statistical analyses since our dataset contained animals from two separate studies but future studies could determine if the difference suggested by Figure 6 is replicable.

**Figure 6.**
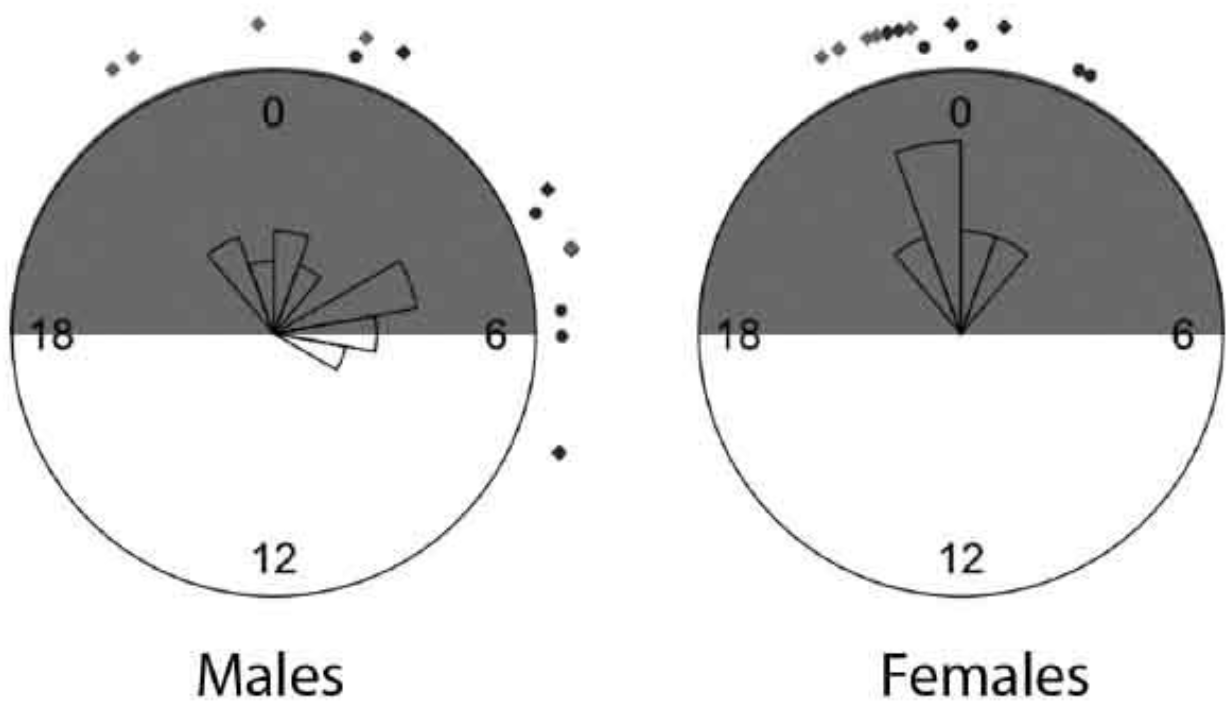
DWT peak circular plots. Plots show DWT peaks near day 2 mark., 0=midnight; mice came from a colony room with lights on 6am-6pm. Shaded grey area indicates lights off in the colony room. DWT peak marker times for data from mice with osmotic mini pumps. Circles placed in inner ring = 7-day pump, diamonds placed in outer ring = 14-day pumps, dark grey = mPer2^Luc/Luc^ mice included in this study, light grey = m*Per2 ^LucSV/+^* controls (No Cre Bmal1^fl/fl^) from previous study (van der Vinne et al. 2020). Jitter (factor = 3) was added to the images to reveal overlapping data points.

We were not able to find any studies describing sex differences in whole-body PER2::LUC bioluminescence. Some sex differences in *in vitro* bioluminescence of explant tissues have been studied. Amplitude of the PER2::LUC rhythm in the adrenal and liver is higher in male mice (Kuljis et al. 2013, Kloehn et al. 2016), and phase of liver and adrenal rhythms was phase advanced in male mice (Kuljis et al. 2013). *In vitro* retinal PER2::LUC oscillations show no sex-dependent effects on amplitude, period, phase, or rhythmic power (Calligaro et al. 2020). Sex differences in sleep and circadian variables may play an important role in application of circadian research on negative health impact of circadian disruption (Boivin et al. 2016, Yan and Silver 2016, Qian et al. 2019).

### Limitations

We have discovered several methodological concerns. Due to the impact of room temperature fluctuations on PMT noise, our technical approach requires either carefully controlled room temperature or (ideally) a method for correction of temperature-related noise on the PMTs. We solved this problem by collecting dark counts regularly and subtracting those measures. Cooled PMTs might offer further advantages.

We initially attempted to study mice with E-mitter implantable telemetry probes (Starr Life Sciences) to monitor rhythms in body temperature simultaneously with bioluminescence. Unfortunately, when implanted in a mouse housed in a LumiCycle *In Vivo* unit, the telemetry probes created electronic noise that appeared as additional PMT signal especially when the mouse showed high levels of locomotor activity. We confirmed this conjecture by detecting rhythmic “bioluminescence” from a mouse with an implanted telemetry probe, but with no substrate delivered by pump or drinking water (data not shown). A different telemetry system might offer the ability to monitor body temperature and other rhythms.

While the body temperature probes could not be used as intended, their presence in animals recovering from the subsequent osmotic pump implant surgery provided insight into the prolonged effects of general anesthesia in our mice. During recovery from surgery, we noted that the circadian rhythm in body temperature was disrupted for 1-2 days, similar to prior reports but for a longer duration (Shirey et al. 2015). Based on these results, we allowed 3 days for colony room recovery after osmotic pump implantation surgery in the studies reported here. Alterations in the circadian rhythms in body temperature might interfere with the hypothesized role of body temperature rhythms to synchronize rhythms in gene expression in peripheral tissues (Brown et al. 2002, Buhr et al. 2010, Schibler et al. 2015). The 3-day recovery period is a significant portion of the period of pump life from the 7-day, emphasizing the need to find alternative routes of administration or substrates that can be more concentrated to accommodate the fixed minipump volume, thus allowing longer studies. Our use of drinking water to deliver luciferin allows us to avoid the use of anesthesia and surgery, thus avoiding these potential effects on body temperature rhythms.

Cycles in skin pigmentation and hair regrowth after shaving of pigmented animals can interfere with bioluminescence imaging (Curtis et al. 2011) and should be considered when planning longer-term data collection. At the time of shaving, we noted variations in skin color and texture. Some animals exhibited a mottled skin color on both back and abdomen, with irregular patches of dark pigmentation. These variations likely arose from the stage of hair growth the mouse was in at the time of shaving, which does not take place uniformly across the animal but rather occurs in waves (Plikus and Chuong 2008). In some cases hair removal itself can cause more rapid hair regrowth, or the animal may have already been in an active stage of the hair regrowth cycle (Li et al. 2017). Future studies to address this may include crossing transgenes onto an albino background to avoid the pigmentation and possibly even avoid the need for shaving. A hairless mouse might provide optimal background (Collaco and Geusz 2003, Izumi et al. 2017) although hairless mice might still have skin pigmentation and can show other alterations in their phenotype (Hoshino et al. 2017). Moreover, there are sex differences in the optical properties of murine skin to be considered in experimental design. Males show an increase in reflectance and scattering of the bioluminescent signal due to their generally thicker dermis (Calabro et al. 2011). Consequently, careful consideration of timing of shaving as well as hair regrowth patterns must be taken to ensure optimal bioluminescence recordings. As described, the variability of pigmentation and hair regrowth patterns might be very limiting when recording from a tissue-specific area. Future studies to optimize *in vivo* bioluminescence imaging may include crossing transgenes onto an albino background to avoid pigmentation. Both albino and hairless phenotypes arise from recessive alleles; generating reporter mice with either albino coat or hairlessness requires at least 2 generations of breeding and might preclude generating more complicated genetic combinations (Konger et al. 2016).

### Future directions

We anticipate further developments that will continue to optimize this approach for study of the circadian system. We look forward to studies of mice with various bioluminescent reporters, other synthetic substrates, and varied delivery methods. It is key to develop tissue-specific circadian reporters, and substrates may then need to be developed with each target tissue in mind. Further innovation is possible as this method expands. We did not test other PMTs available but innovations here might allow better detection of signals. Note that although firefly luciferase emits green light at room temperature, this emission is red-shifted at 37 °C and further red-shifted by depth-dependent spectral differences in transmission through the body (Zhao et al. 2005). PMTs with increased sensitivity at long wavelengths may thus be desirable. We expect that further optimization of these methods is possible, but here we demonstrate a method that reliably detects circadian rhythms in bioluminescence from the mPer2^Luc^ mouse.

## Acknowledgements

This work would not have been possible without the technical assistance and innovation of Dr. David Ferster. We would like to thank Donna Mosley and Juliane Donahue Bombosch for assistance with preparation of figures. We thank Halley Lin-Jones and Melissa Chenok for helpful discussions. This research was supported in part by NIH R15GM126545 to MEH, NIH R21NS103190 to DRW, and R01EB013270 to SCM. The content is solely the responsibility of the authors and does not necessarily represent the official views of the National Institutes of Health.

## Editorial Note

Dr. William J. Schwartz, previous editor of the *Journal of Biological Rhythms* graciously agreed to handle the editorial process surrounding review of this manuscript. Neither the current Editor of the *Journal* (MEH) or the Deputy Editor (DRW) had any role in the review process and will remain blind to the reviewers’ identities.

## Supplementary Methods

### Animal Housing

Standard mouse cages measuring 7 1/2” × 11 1/2” × 5” (Ancare, Bellmore, NY) with 30 g TekFresh bedding were used for all experiments in the LumiCycle *In Vivo* units, with a few simple modifications. To minimize obstruction in the field of view of the PMTs, cage lids were modified by removing the divider that normally contains chow. Chow (Teklad 2014) was available in a stainless-steel hopper (Ancare) side-mounted inside the cage. An aluminum bar measuring 4” × 1” × 5/8” was placed under the hopper to keep mice from nesting beneath. Reduced height square water bottles (Ancare) with #6 stoppers allowed the cages to be moved easily in and out of the unit. For delivery of a substrate in drinking water, 50 mL conical tubes were used instead of water bottles, also with # 6 stoppers. To ensure the availability of the solution through the stopper, the angle of the tubes was kept constant by an extension spring clipped to the cage lid (See Supplementary Figure 2). We recorded general locomotor activity by a passive infrared motion sensor using Clocklab software (RRID:SCR_014309).

Possible sources of background noise include the type of bedding and the composition of the diet. To rule out these possible contributors in our system we tested an empty cage as well as ones bedded with Aspen bedding, white crinkled paper, and two amounts of TekFresh (7099, Envigo). The lesser amount of TekFresh was closest to the level of the empty cage and all subsequent experiments used that amount. To avoid abdominal autofluorescence as reported (Inoue et al. 2008) we used a global, alfalfa-free diet (Teklad 2014, Envigo). We did a further comparison between that and a purified diet (AIN-93M, Envigo) and found no significant difference between the two in our system.

### Pump Preparation

Alzet^®^ subcutaneous osmotic minipumps (Durect, Cupertino, CA), calibrated to deliver for either 7 or 14 days were used. On the day prior to implantation, pumps were filled with substrate which had been sterile-filtered using 1 mL Luer Slip syringes (Grainger Industrial Supply, Springfield, MA) and Millipore syringe filters (Thermo Fisher Scientific, Waltham, MA). 0.5cc BD Insulin Syringes with no residual volume (MWI Veterinary Supply, Boise, ID) were used to fill the pumps. To measure the final volume in the filled pump, each pump, flow moderator, and packaging was weighed before and after filling. Weights were recorded to determine initial filled volume and to compare with the residual volume at the end of treatment, to measure the actual delivered dose. Pumps were then primed for up to 24 hours in 1.5 mL sterile saline in 15 mL conical tubes, at 37° C.

### Surgery for Pump Implantation

One hour prior to surgery animals were weighed and injected subcutaneously with buprenorphine (0.05 mg/kg) and either ketoprofen (5 mg/kg) or meloxicam (2 mg/kg) as analgesics. At the beginning of surgery animals were placed in an induction chamber with 3% isoflurane, then transferred to a nose cone. Body temperature was maintained with a heated gel pack placed under the animal throughout surgery. Eyes were protected with veterinary ophthalmic ointment and anesthesia was maintained at 2.5% to 3%.

Animals were shaved with an OsterFinisher clipper (Osterpro.com) on the back, from shoulder blades to pelvic bones, and around both sides and the complete abdomen (See Supplementary Figure 3). After washes with Betadine and 70% EtOH and using sterile surgical technique, a ~10 mm skin incision with a #15 scalpel was made below the shoulder blade, perpendicular to the spine. A 20 mm subcutaneous pocket was created using a blunt hemostat and flushed with 0.5 mL sterile saline. Pumps were removed from the priming solution and inserted into the pocket. Incision was closed with 3-4 absorbable sutures (Ethicon #023434, MWI Veterinary Supply, Boise ID) and treated with a splash block (Lidocaine 0.2%, Bupivicaine 0.05%) and antibacterial ointment (Vetropolycin, MWI Veterinary Supply).

During recovery animals were housed individually in large standard mouse cages (10 1/2” × 19” × 6 1/8”, fitted with Micro Filter Top™, Ancare) with TekFresh bedding. Cages were placed on top of heated discs (SnuggleSafe.com) overnight and animals monitored for recovery and freedom of movement. Besides normal chow ad libitum and water, soft food and apple slices were placed in dishes on the cage floor. After three days recovery in 12:12 LD, animals were moved into smaller standard cages and placed into individual LumiCycle *In Vivo* units in constant darkness, just prior to the time of lights off in 12:12 LD. Data were recorded for 7 days in DD, and animals were checked once a day at randomized times using an infrared viewer. As well as bioluminescence, locomotor activity was recorded throughout the experiment with a passive infrared motion sensor using ClockLab (Actimetrics). After completing the pump experiments, animals were euthanized and tissue samples were collected for confirmation of genotype.

### Surgery for Pump Removal

For those experiments in which an animal was to be tested with a second pump after recovery, a pump removal protocol was followed. Animals were briefly anesthetized with 3% isoflurane in an induction chamber, to allow for subsequent subcutaneous injections of buprenorphine and ketamine or meloxicam as previously described, without restraint. Animals were returned to their home cages for at least 30 minutes before beginning surgery. An additional brief induction followed by transfer to a nose cone preceded the pump removal and body heat was again maintained with a gelpack. Any remaining sutures were removed and an incision was made in the original site or parallel to it, depending on the degree of healing present. The pump was removed, the subcutaneous pocket flushed with 0.5 mL saline and incision closed as for pump insertion. Postoperative care was the same as for pump insertion.

### Dose determination

Immediately following a pump removal, any remaining substrate was removed from the pump using a 0.5 cc BD Insulin Syringe. Actual dose delivered was calculated by the following formula, as provided by the manufacturer Alzet:

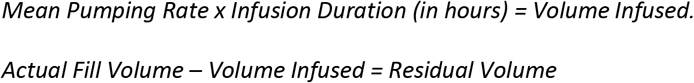

During the hours of priming before surgical implantation in the animals, the rate per hour was calculated at half the mean pumping rate. From the time of implant until removal the rate was calculated at the normal pumping rate.

### Thermal stability of luciferins

D-luciferin sodium salt was purchased from GoldBio. CycLuc1 was synthesized as previously described (Reddy et al. 2010). The luciferin substrates were dissolved in PBS to a final concentration of 100 mM (D-luciferin) and 5 mM (CycLuc1), then incubated at 4 °C or 37 °C for the indicated time (0-14 days). Aliquots were removed for LC-MS and bioluminescence imaging comparisons.

FVB/NJ mice were purchased from Jackson Laboratories (Bar Harbor, ME). Intravenous injection of AAV9-CMV-WTluc2 was performed as previously described (Mofford et al. 2015). Bioluminescence assays were performed on a Xenogen IVIS-100 system in the UMass Medical School Small Animal Imaging facility. All luciferin aliquots were sterile-filtered through a 0.22 μm syringe filter (Millex-GV) prior to injection. Each mouse was weighed to determine substrate dosing and anesthetized using 2.5% isoflurane in 1 L/min oxygen. Each luciferin substrate was injected i.p. at a dose of 4 μL/g mouse and mice were imaged ventrally 12 minutes after injection. Data acquisition and analysis were performed with Living Image^®^ software. Data were plotted and analyzed with GraphPad Prism 7, and reported as the total flux (p/s) for the whole mouse. All these experiments were conducted in accordance with the Institutional Animal Care and Use Committee of The University of Massachusetts Medical School (docket #A-2474-14).

LC-MS analyses were performed on an Agilent Technologies 6130 quadrupole LC-MS connected to an Agilent diode array detector. Chromatography was conducted on an Agilent InfinityLab Poroshell 120 EC-C18 column (4.6 × 50 mm, 2.7 micron particle size), with a mobile phase of water/acetonitrile with 0.1% formic acid. Samples were prepared by dilution of an aliquot of the 100 mM and 5 mM PBS stocks with Milli-Q water to 500 μM D-luciferin and 250 μM CycLuc1, respectively. Of this solution, 20 μL was injected onto the C18 column, and a gradient of H2O and acetonitrile (+0.1% formic acid) was used as the mobile phase.

| Time (min) | Water (0.1% Formic acid) | Acetonitrile (0.1% Formic acid) | Flow rate ml/min | Pressure (bar) |
| --- | --- | --- | --- | --- |
| 2 | 100 | 0 | 1.0 | 600 |
| 10 | 0 | 100 | 1.0 | 600 |
| 12 | 100 | 0 | 1.0 | 600 |
| 14 | 100 | 0 | 1.0 | 600 |

### Materials

| **Name** | **Company** | **Catalog Number** |
| --- | --- | --- |
| D-Luciferin Potassium Salt | GoldBio | Catalog # LUCK-1G |
| CycLuc1 | gift of Dr. Steve Miller |  |
| Alzet osmotic minipumps 7 day | Durect | Model 1007D Cat # 0000290 |
| Alzet osmotic minipumps 14 day | Durect | Model 1002 Cat #0004317 |
| Luer slip syringes | Grainger Industrial Supply | Item # 19G334 Mfr. Model # 4010.200V0 |
| Millipore syringe filters | Thermo Fisher Scientific | CAT # SLGV004SL |
| 0.5cc BD Insulin Syringes | MWI Veterinary Supply | MWI SKU: 19545 |
| Vetropolycin Antibiotic Ointment | MWI Veterinary Supply | MWI SKU: 001479 |
| Oster Finisher clipper | OsterPro.com | Oster® Finisher® Trimmer (Model 59) |
| Ethicon Absorbable sutures | MWI Veterinary Supply | Cat # 023434 |
| Recovery cage | Ancare | N40 |
| Recovery cage heat discs | SnuggleSafe | Microwave heat pad |
| Micro Filter Top | Ancare | N40 Micro Filter Top |
| Recording cage | Ancare | N10 |
| Reduced Height Square water bottle | Ancare | Reduced Height Square Bottle |
| Food hopper | Ancare | Mouse Feeder |
| Carson OPMOD DNV viewer | www.opticsplanet.com | 99-NV-OPMOD-DNV-DN-300 |
| Pulsar Edge Night Vision Goggles | www.nightgearstore.com | PU-75095 |

### Supplementary Figure captions

**Figure S1.**
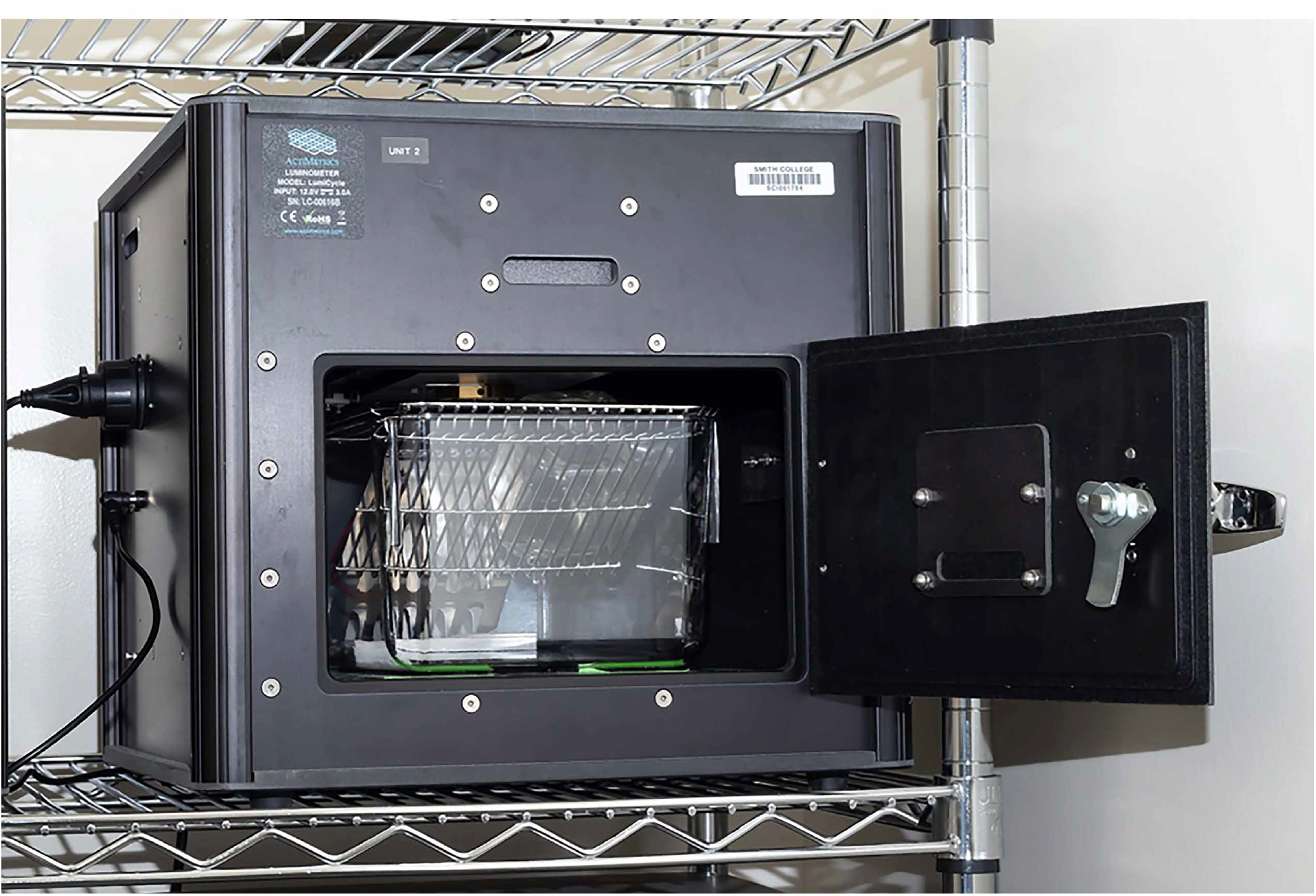
The LumiCycle *In Vivo* System, single unit

**Figure S2.**
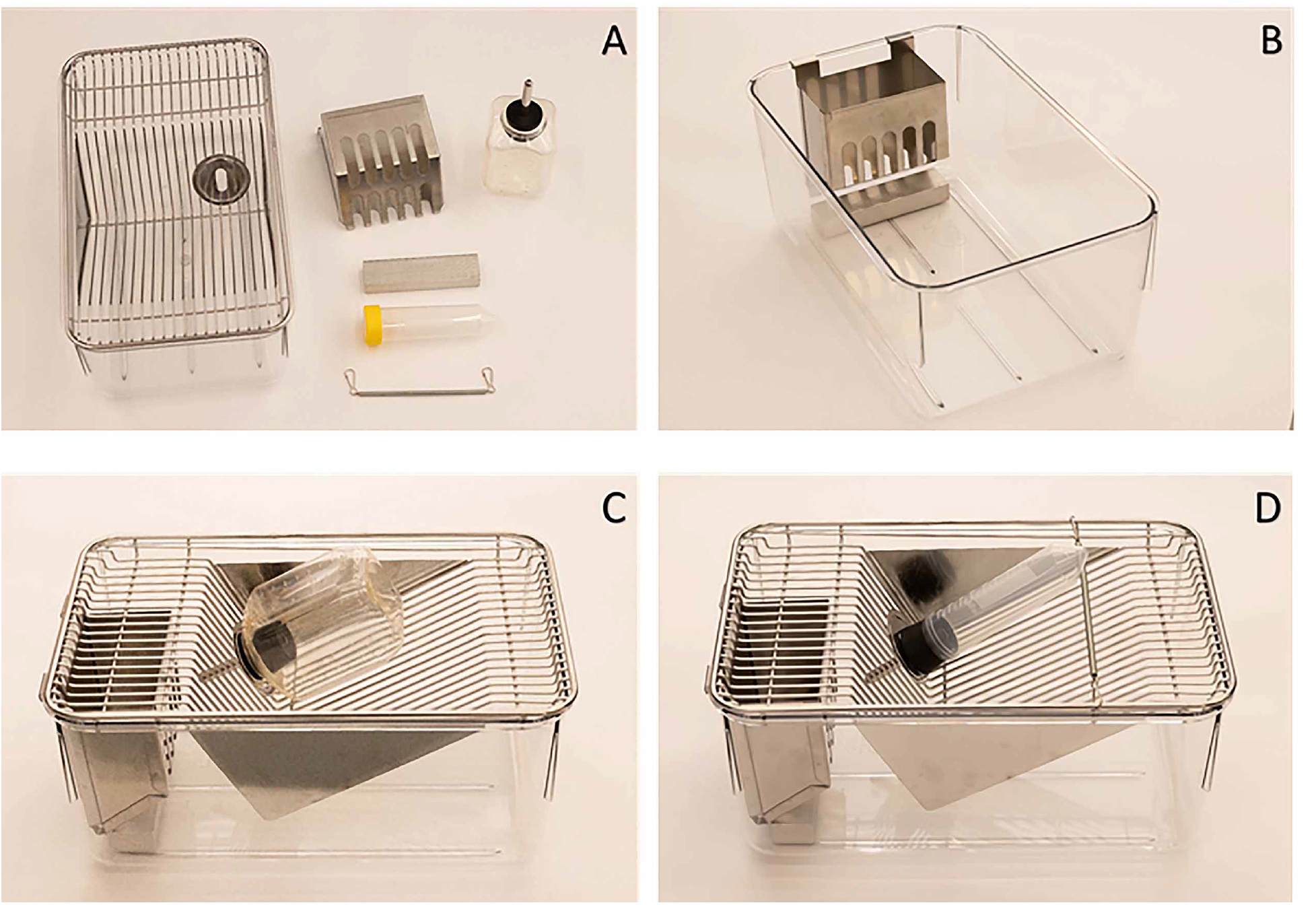
Standard cage and accessories (A) Standard mouse cage, food hopper, aluminum bar, reduced height water bottle or conical tube and extension spring to maintain angle for small volume solutions. (B) Aluminum bar is placed underneath food hopper to prevent nesting and to maximize visible range of PMTs. (C) Cage assembly with reduced height water bottle. (D) Cage assembly with conical tube and extension spring to maintain angle of tube when using low volumes of substrate solutions.

**Figure S3.**
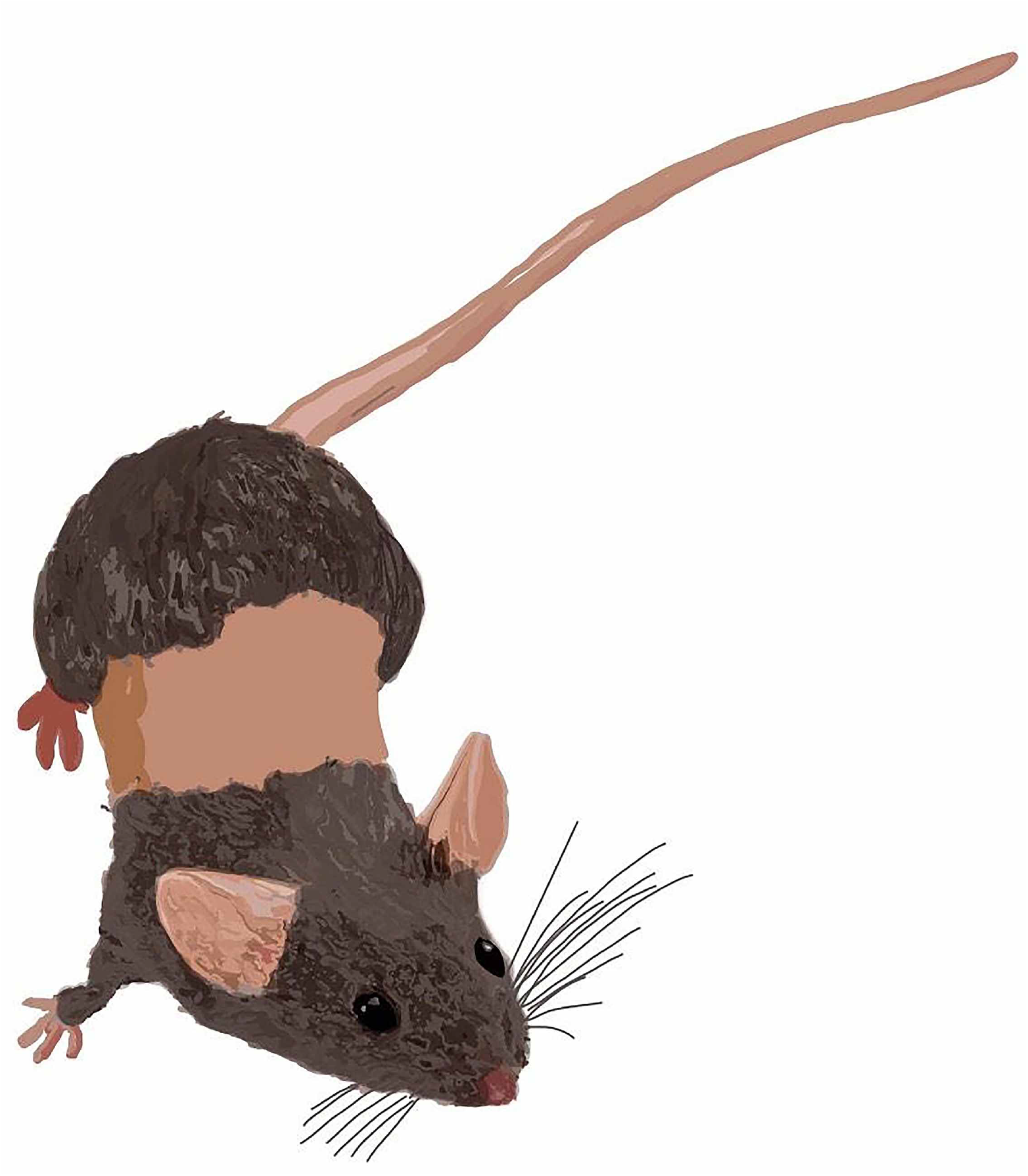
Drawing to demonstrate the extent of the shave administered prior to experiments. The mouse was shaved in a band that extended from dorsal to ventral surface.

**Figure S4.**
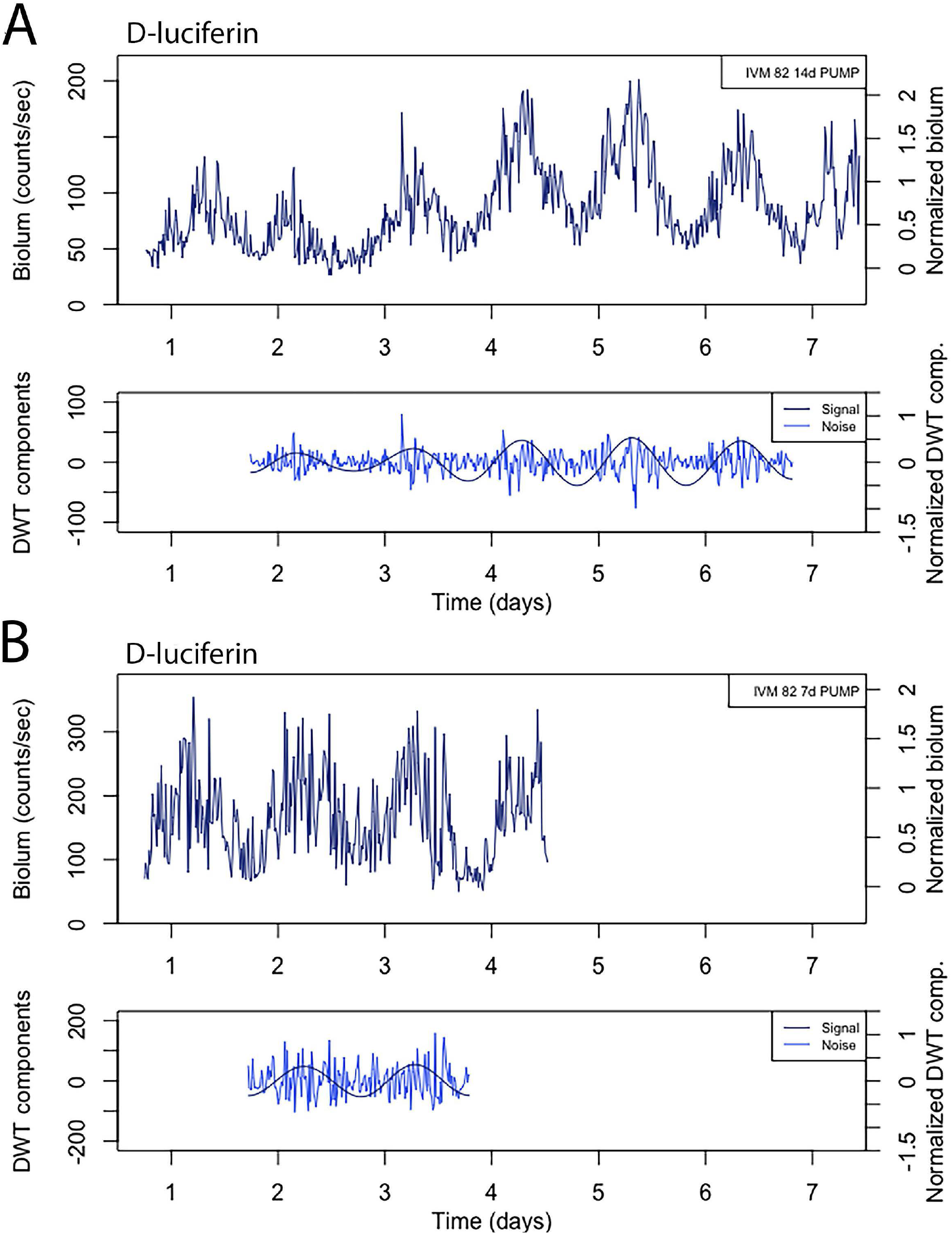
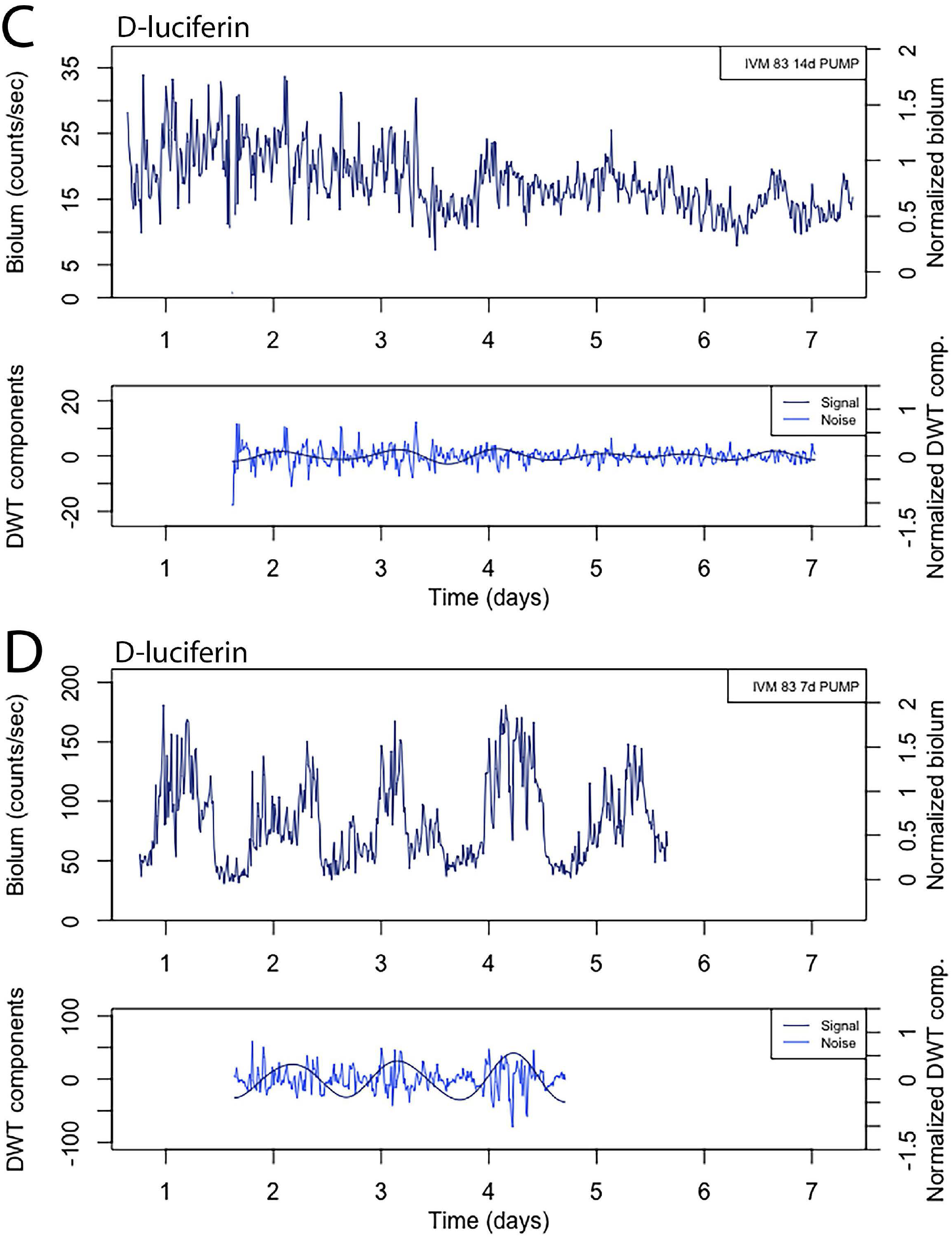
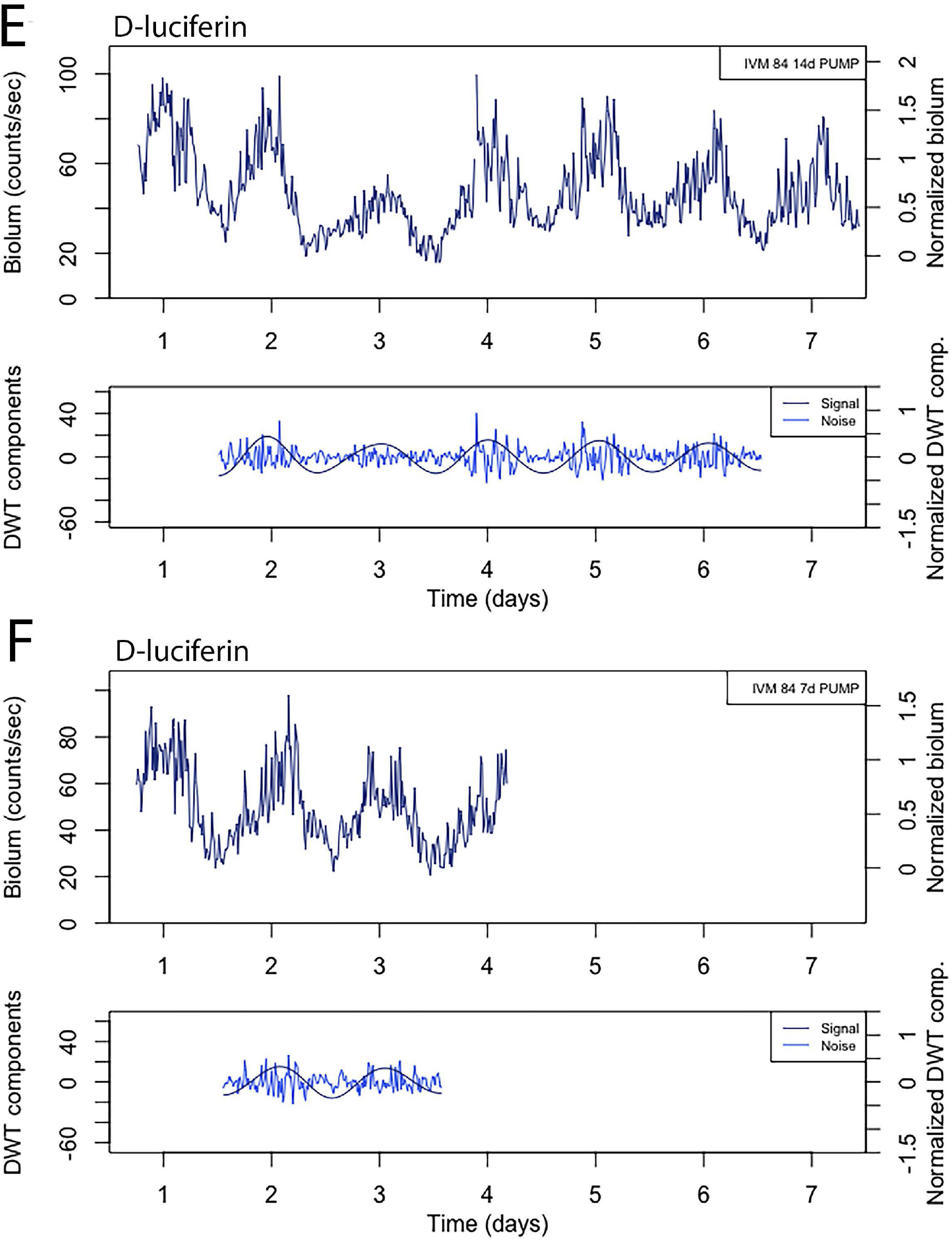
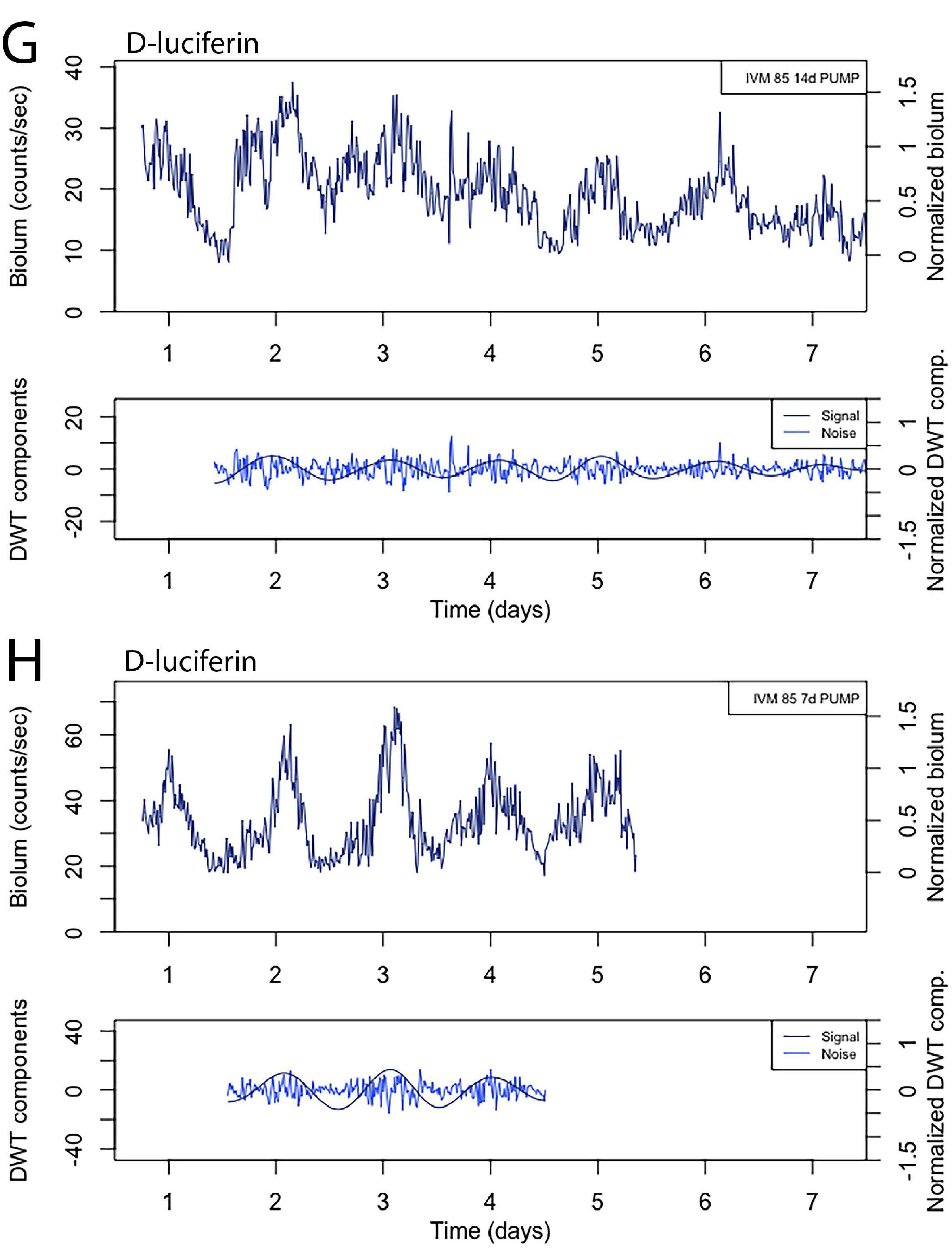
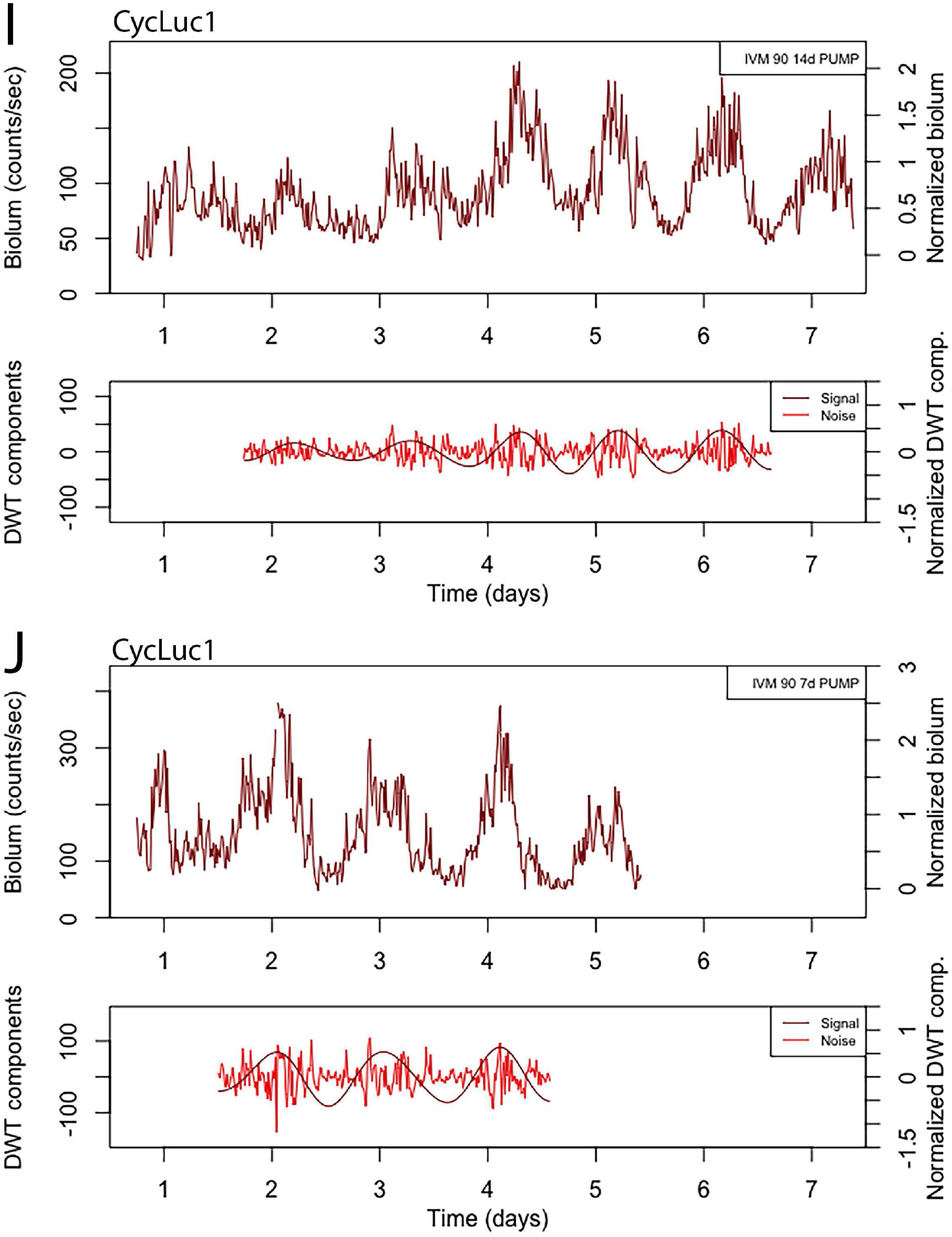
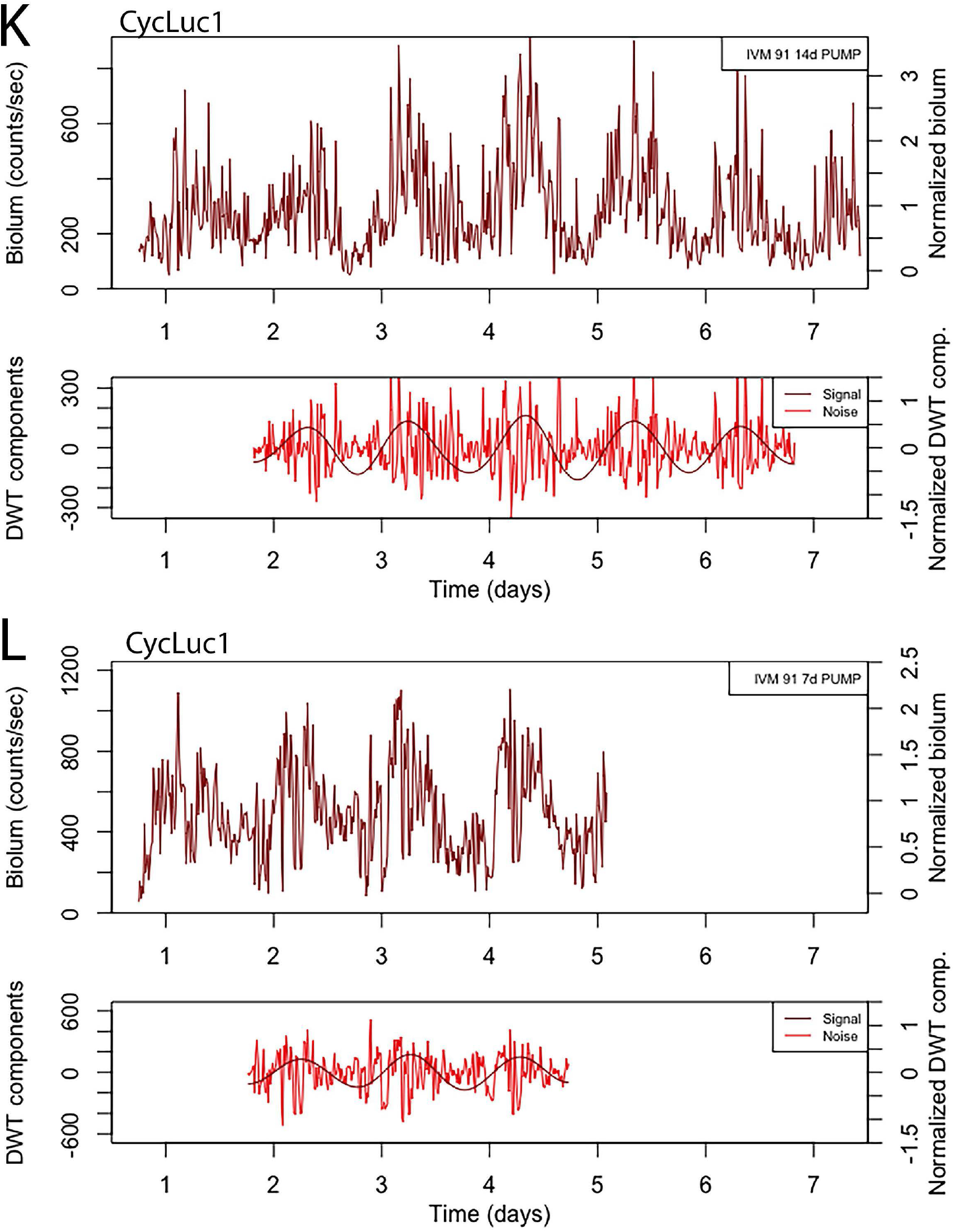
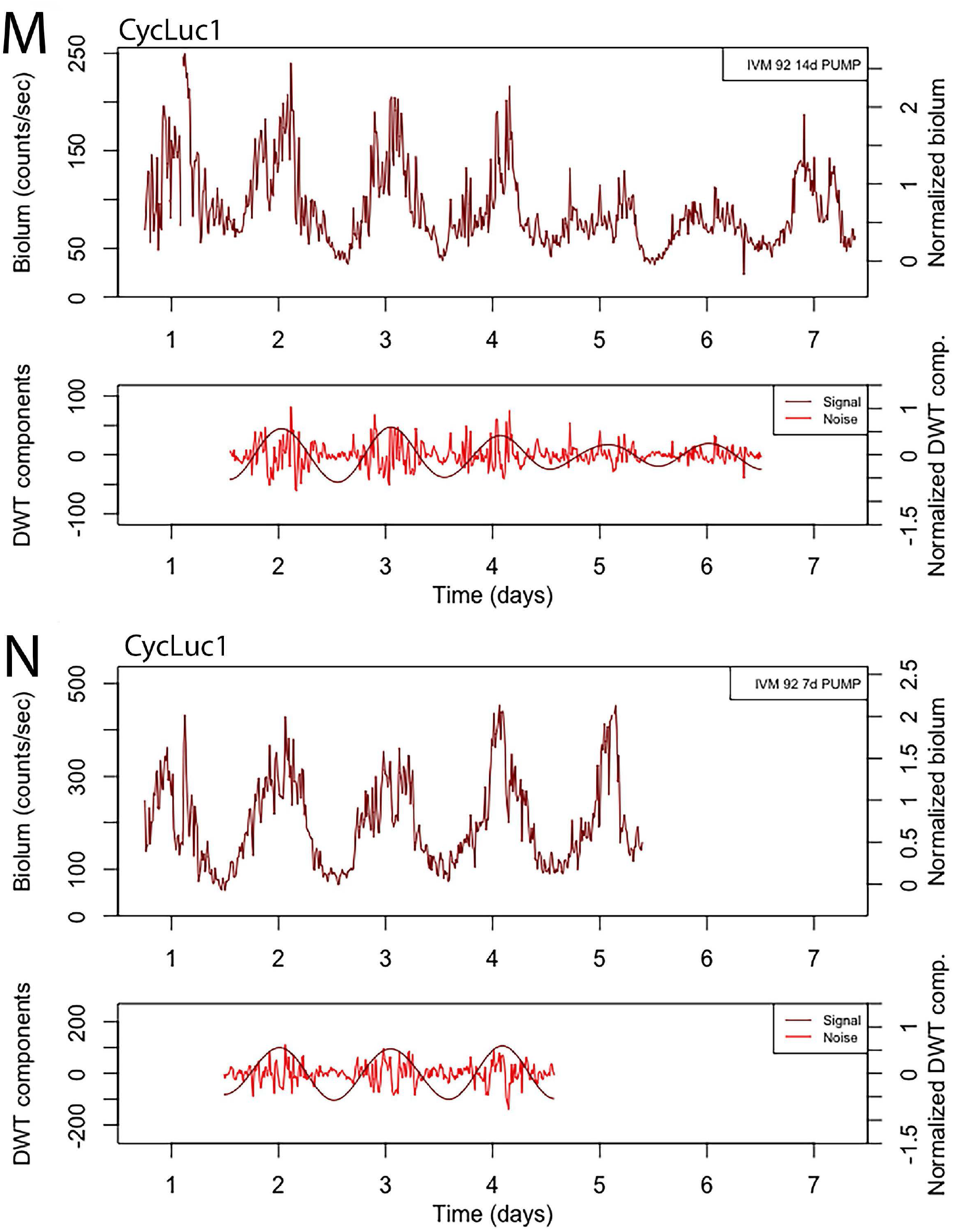
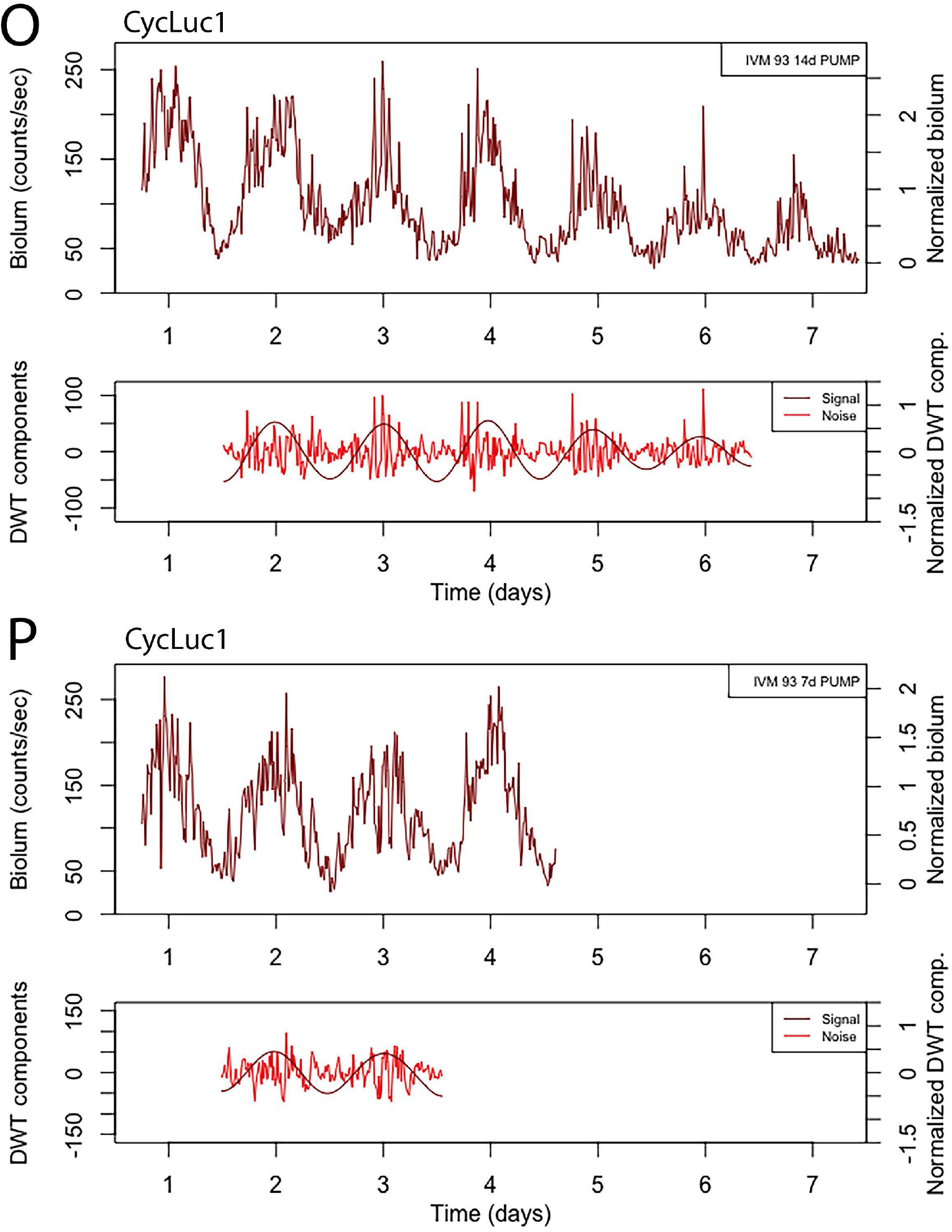
All results of bioluminescence data with either 7-day or 14-day D-luciferin or CycLuc1 pumps. Top graph for each example: 15-minute median binned trace with counts/sec scale on the left and normalized scale on the right (subtract first percentile then divide by median, so min is mapped to zero and median mapped to 1; using first percentile reduces the effect of outliers). Bottom graph for each example: For the signal-to-noise ratio analysis, the DWT-calculated circadian component D6 is treated as the signal and the summed components D1-D4 are treated as the noise. The data before the first trough and after the last trough are discarded to avoid edge effects. The 14 day pumps delivered 0.25 uL/h of 100 mM D-Luciferin or 5 mM CycLuc1. The 7 day pumps delivered 0.50 uL/h of 100 mM D-Luciferin or 5 mM CycLuc1 (blue - D-luciferin pumps, red-CycLuc1 pumps). Individual animals are identified by the “IVMxx” number.

**Figure S5.**
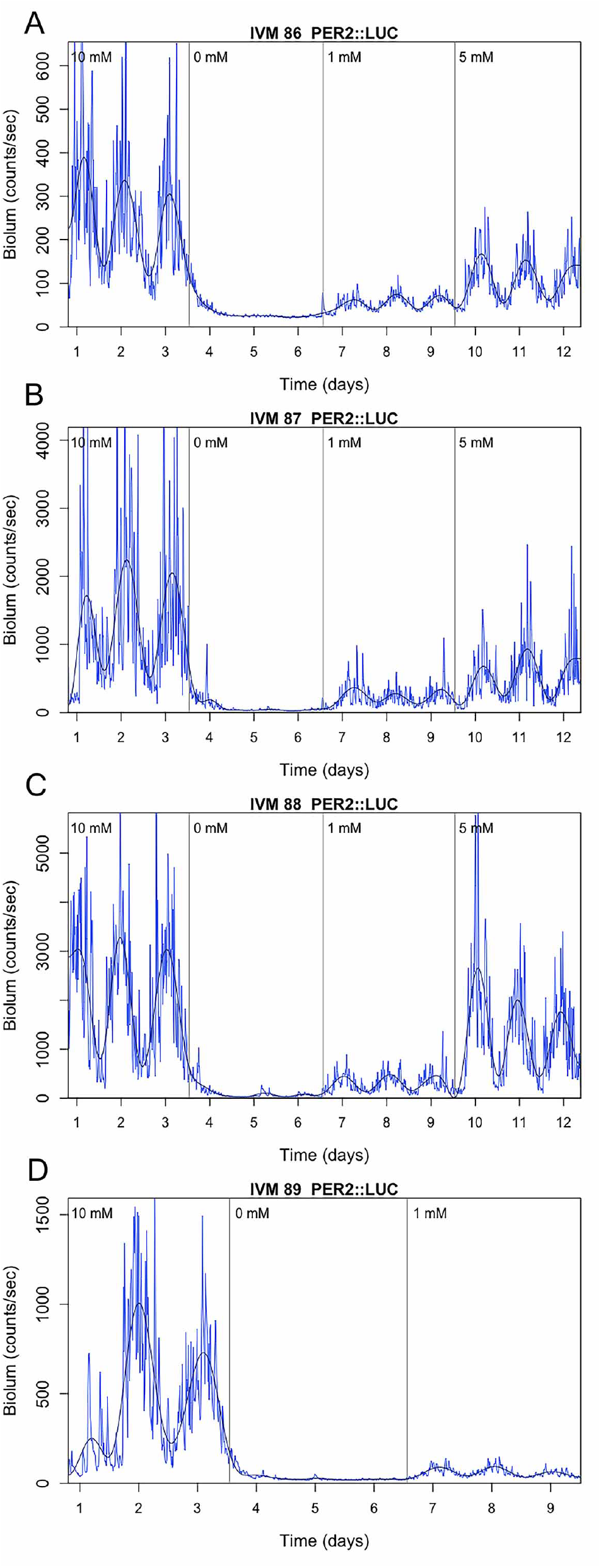
All drinking dose response data. Individual animals are identified by the “IVMxx” number. Final data for IVM 89 (D) was lost due to equipment failure.

**Figure S6.**
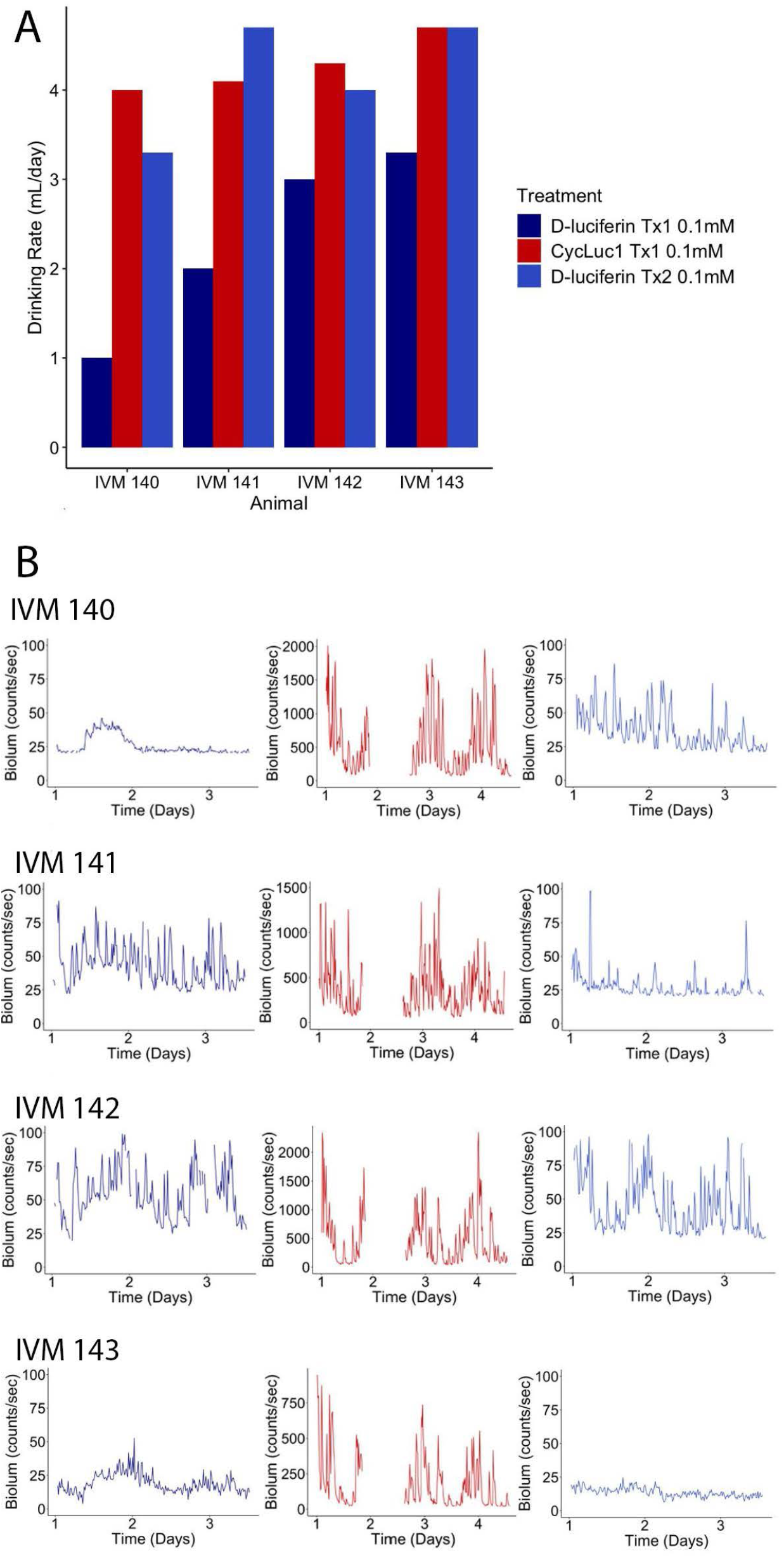
A) Drinking rate results for CycLuc1 vs D-luciferin comparison data. Drinking rate per day for each treatment was calculated by dividing the total volume of substrate consumed by the number of days. B) Bioluminescence records for all animals (Left - 0.1 mM D-Luciferin treatment 1, center - 0.1 mM CycLuc1 treatment, right - 0.1 mM D-Luciferin treatment 2). Individual animals are identified by the “IVMxx” number.

**Figure S7.**
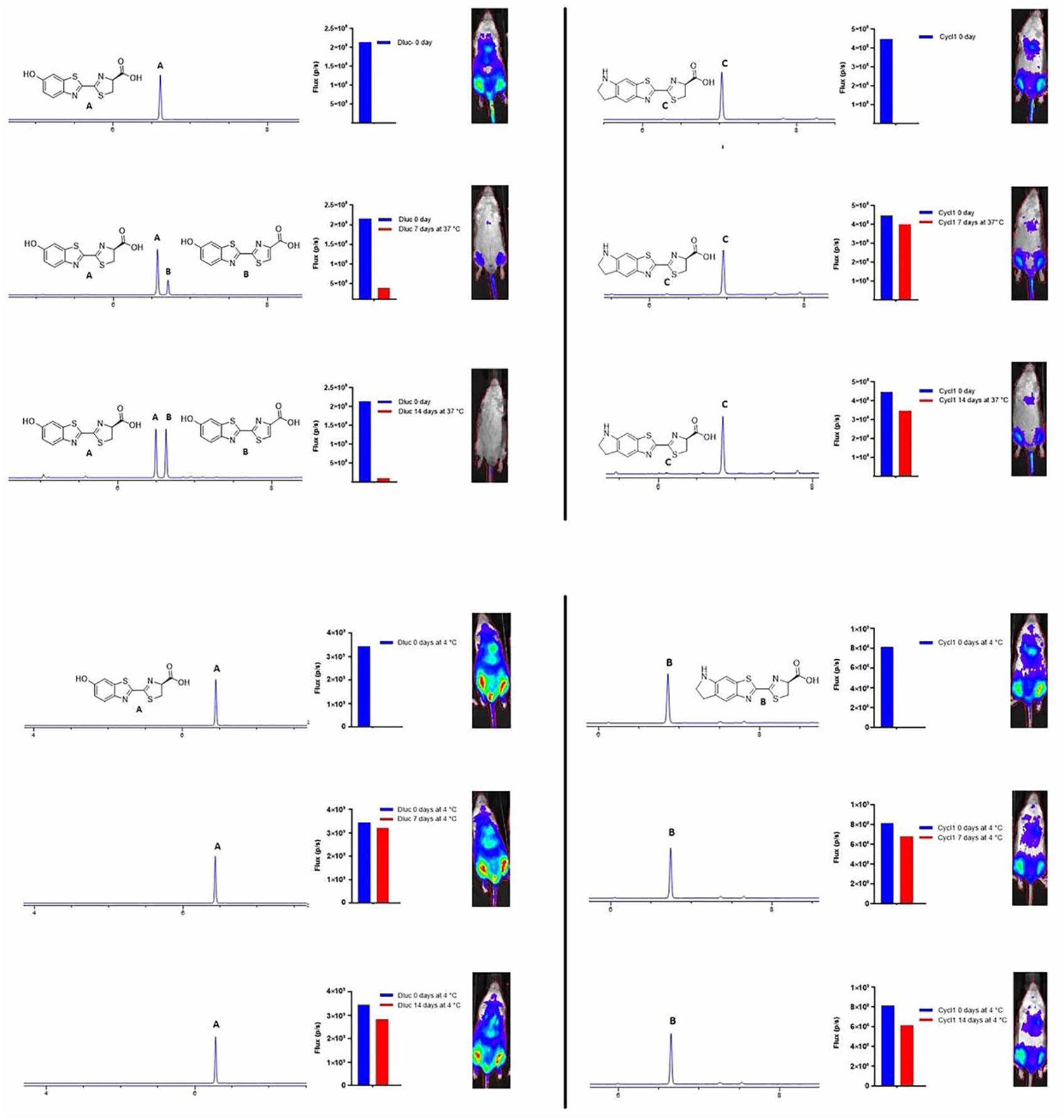
Evaluation of D-luciferin and CycLuc1 after incubation at 4 °C or 37 °C. LC/MS and in vivo bioluminescence imaging after incubation in aqueous buffer at the specified temperature for 0, 7, or 14 days in FVB/NJ mice previously receiving iv injection of AAV9-CMV-WTluc2 as previously described (Mofford et al. 2015).

